# A multi-omic, spatial, and whole-slide image dataset of lung neuroendocrine tumours from the lungNENomics cohort

**DOI:** 10.64898/2026.05.12.724489

**Authors:** Lipika Kalson, Alexandra Sexton-Oates, Emilie Mathian, Catherine Voegele, Alex Di Genova, Zhaozhi Li, Jaehee Kim, Leigh M. Marsh, Luka Brcic, Lynnette Fernandez-Cuesta, Matthieu Foll, Nicolas Alcala

**Affiliations:** Otto Loewi Research Center, Lung Research Cluster, Medical University of Graz, Graz, Austria; Computational Cancer Genomics Team (CCG), Genomic Epidemiology Branch (GEM), International Agency for Research on Cancer/World Health Organisation (IARC/WHO), Lyon, France; Instituto de Ciencias de la Ingeniería, Universidad de O’Higgins, Rancagua, Chile; Centro UOH de Bioingenieria (CUBI), Universidad de O’Higgins, Rancagua, Chile; Centro de Modelamiento Matemático UMI-CNRS 2807, Santiago, Chile; Department of Computational Biology, Cornell University, Ithaca, USA; Department of Pathology, Hospital Graz II, Graz, Austria l Department of Pathology; Medical University of Vienna, Vienna, Austria

**Author notes:** co-supervision (Matthieu Foll, Nicolas Alcala).

## Abstract

Lung neuroendocrine tumours (lung NETs) are rare neoplasms comprising approximately 2% of lung cancers. Recent studies have identified distinct molecular groups based on transcriptome and methylome data, but genomic and morphological features remain underexplored due to limited whole-genome and imaging data. We have generated the largest multi-omic dataset of lung NETs to date (201 participants, for a total of *n* = 294 tumours), including RNA sequencing, EPIC 850K methylation arrays, and whole-genome sequencing. This multiomic dataset also include multi-regional whole-genome sequencing for 41 participants, allowing for the quantification of intra-tumoural heterogeneity. We additionally generated spatial proteomics (64 participants), spatial transcriptomics (4 participants) and whole-slide histopathology images for 212 cases. This dataset enables a comprehensive characterization of lung NET molecular groups and the identification of group-specific morphological features using deep learning algorithms. All quality control analyses, processed data, and scripts are provided to ensure reproducibility. This dataset is available as a basis for further molecular and morphological analysis of lung NETs, and for future research on multi-scale integration.

## Background & Summary

Lung neuroendocrine tumours (lung NETs) are a rare and understudied type of lung cancer believed to originate from pulmonary neuroendocrine cells^1^, with increasing incidence and limited dedicated therapeutic options^2^. Lung NETs are classified into grades 1 and 2, also known as typical and atypical carcinoids, respectively, based on morphological criteria, including mitotic count and presence or absence of necrosis^3,4^. Although most lung NETs progress slowly and have a good prognosis, some patients, particularly those with grade-2 tumours, do experience relapse and metastases^5^. The current morphological classification is subject to considerable inter-observer variability, hampering its prognostic value, and has reduced therapeutic value^6,7^.

To address these gaps, we established the lungNENomics cohort, a multi-modal dataset providing an integrated molecular and morphological reference for lung NETs^8,9^. Omics studies have previously revealed multiple molecular groups using mainly transcriptomic data; using the lungNENomics dataset presented here, through integration of whole-genome sequencing (WGS), RNA-sequencing (RNA-seq), and DNA methylation arrays, we have shown that lung NETs can be classified into four groups, Ca A1, Ca A2, Ca B, and supra-carcinoid enriched, each with distinct molecular profiles and morphological features^10^.

The dataset includes WGS, RNA-seq, and DNA methylation arrays, spatial data (GeoMx® Digital Spatial Profiler spatial proteomics and 10x Genomics Visium transcriptomics), and high-resolution whole-slide images (WSI) of tumour tissues from the lungNENomics project for the characterisation of lung NETs^8^. The datasets contain molecular data for 201 participants comprising 294 tumour samples in total, including multi-regional samples from 41 participants. The cohort includes *n*=104 grade-1 and *n*=40 grade-2 tumours. Additionally, histopathological image data are available for *n*=212 participants. Along with raw and processed data, we provide quality controls for each technique and scripts to run a complete molecular analysis. All scripts are publicly available on GitHub^11^.

This dataset is the largest multi-omic dataset of lung NETs, particularly valuable given the disease’s rarity and the current paucity of sequencing and imaging data production. We believe this unique dataset will serve as a reference for future studies of this understudied tumour type.

## Methods

### Sample Collection

Fresh-frozen and formalin-fixed paraffin-embedded (FFPE) tumour tissues, fresh-frozen adjacent normal lung tissue, whole blood, and haematoxylin/eosin (HE) or haematoxylin/eosin/saffron (HES) stained slides were collected at diagnosis in the context of the lungNENomics project from 12 contributing centres in Europe and Australia (https://rarecancersgenomics.com/lungnenomics/); see ^10^ for a full description of the cohort. Central pathology review by six pathologists was undertaken on HE/HES slides for 187 of the 201 participants.

### DNA & RNA extraction

For each fresh-frozen tumour sample, DNA extraction was performed either using the Gentra Puregene Tissue Kit from Qiagen (between 2018 and 2020), or the DNAdvance Tissue Kit (since 2021), following the manufacturer’s instructions. RNA extraction was performed either using the miRNAeasy Mini Kit (between 2018 and 2020), or the RNAdvance Tissue Kit (since 2021), following the manufacturer’s instructions.

### Sequencing

WGS was performed for 72 participants, all with matched normal tissue or blood, RNA-seq for 178 participants, and DNA methylation arrays for 191 participants. All three omics data types were obtained for 60 participants, while 109 had both RNA-seq and DNA methylation array data. For 41 participants, multiple regions (between two and seven) from the same tumour were sequenced. Spatial transcriptomics was performed for four participants, and spatial proteomics was performed on 64 participants.

#### WGS

WGS was performed by the Centre National de Recherche en Génomique Humaine (CNRGH, Institut de Biologie François Jacob, Commissariat à l’énergie atomique et aux énergies alternatives), France, on 106 fresh-frozen tumours (including 34 multi-region tumour samples) from 72 participants and 72 matched normal tissue or blood samples. The Illumina TruSeq DNA PCR-Free library preparation kit (20015963; Illumina) was used for library preparation, and sequencing was performed on an Illumina HiSeqX5 platform at a target depth of 60x for tumour tissue and 30x for normal samples, as paired-end 150 bp reads.

#### RNA-seq

RNA-seq was performed by the Cologne Center for Genomics, Germany, on 239 tumours (including 61 multi-region samples) from 178 participants. After RNA quality control, libraries were prepared using the Illumina TruSeq Stranded mRNA polyA Kit (20020595; Illumina). Libraries were sequenced using an Illumina NovaSeq 6000, as paired-end 100 bp reads.

#### DNA methylation

DNA methylation arrays were performed at the International Agency for Research on Cancer, France, on 277 tumours (including 86 multi-region tumour samples) from 191 participants. DNA was bisulphite converted using the Zymo EZ-96 DNA Methylation kit and hybridised to Infinium MethylationEPIC v1.0 BeadChip arrays (WG-317-1003, Illumina). Arrays were scanned using the Illumina iScan, generating raw intensity (IDAT) files.

#### Spatial transcriptomics

Spatial transcriptomics sequencing was performed at the Centre Léon Bérard, France, on four tumours from four participants. Material from FFPE blocks was placed on a 10x Genomics Visium v1 slide, then underwent deparaffinisation, HE staining, and decrosslinking steps. Human probes targeting approximately 18,000 genes were hybridised overnight and mRNA captured on each spot was retrotranscribed and amplified following 10x Genomics’ protocols. GEX libraries were prepared and sequenced on an Illumina NovaSeq machine with a target sequencing depth of 50,000 reads per spot.

#### Spatial Proteomics

Spatial proteomics, through digital spatial profiling (DSP), was performed at the Centre Léon Bérard, France, on 64 tumours from 64 participants. A total of 513 areas of interest (AOIs) were selected blindly to molecular group and tumour grade. Immune AOIs were defined based on CD45 expression and tumour cell AOIs based on pan-cytokeratin (PanCk) fluorescence following UV illumination of FFPE slides. After selection, the NanoString GeoMx® DSP system was used to quantify the protein expression of 39 proteins (immune cell, immune activation state, and immune cell typing panels) using the NanoString nCounter platform.

### Whole-slide histopathology

The HE/HES-stained slides were scanned at 40x magnification using a Leica Aperio AT2 (Leica Biosystem) to generate WSIs digitalised by the participating centers and stored as high-resolution image files in *svs* format. Samples underwent histopathological review by a panel of six expert pathologists following the 2021 WHO guidelines as described previously^12^.

### Data Processing

Data processing was performed using the bioinformatic workflows from the Computational Cancer Genomics team at the International Agency for Research on Cancer/World Health Organization (https://github.com/IARCbioinfo/), except for WGS whose primary processing was performed at CNRGH, France. We refer readers to previous publications^8,9,13^ from the team for full details and provide a summary of the key steps below. The workflows are written in the Nextflow domain-specific language^14^, and software dependencies are contained in conda environments and containerised with Docker and Singularity (containers available at https://hub.docker.com/ and https://singularity-hub.org/).

#### WGS

Raw reads were mapped to reference genome GRCh38 (with ALT and decoy contigs). The workflow maps reads (software bwa-mem v0.7.15-r1140), marks duplicates, and sorts the reads (software sambamba; v0.6.8-pre1). Quality control metrics were generated using software FastQC (v0.12.1^15^, workflow fastqc-nf) and qualimap (v2.2.2c^16^, workflow qualimap-nf).

Small variants (single nucleotide variants, multi-nucleotide variants, and Indels) were called using GATK4’s Mutect2 method (gatk v4.2.0.0^17,18^, using our workflow *mutect-nf* v2.3). Indels and multi-nucleotide variants were additionally called using Strelka2 (v2.9.10^19^, workflow *strelka2-nf* v1.2a) and were only retained if they were also identified with Mutect2 to reduce false-positive discoveries. Germline variants were called using Strelka2 only (v2.9.10, workflow *strelka2-nf* v1.2a). Resulting variant calling format (VCF) files were normalised using bcftools (v1.10.2^20^, workflow vcf_normalization-nf v1.1) and annotated using ANNOVAR (v2020Jun08, workflow table_annovar-nf v1.1.1).

Somatic and germline structural variants were identified using our workflow sv_somatic_cns-nf (v1.1) which performs consensus calling combining three structural variant callers (DELLY (v0.8.7^21^), Manta (v1.6.0^22^), and SvABA (v1.1.0^23^). A panel of germline structural variants was generated from normal samples, and somatic variants with overlapping breakpoints (within a 100bp region) of germline events that occurred in more than 1% of samples were removed.

Copy-number variants were called using PURPLE (v2.52^24,25^, implemented in our nextflow pipeline *purple-nf* v1.1), using high quality somatic variant calls to refine copy-number estimates. PURPLE was also used to estimate tumour purity, ploidy, whole-genome duplication status, and microsatellite instability. Recurrent broad and focal copy-number alterations were identified using GISTIC2 (v2.0.23) via the aCNViewer^26^ framework.

#### RNA-seq

Raw reads were mapped to reference genome GRCh38 with annotation gencode v33 using our pipeline *RNAseq-nf* (v2.4). This workflow removes adapter sequences (wrapper Trim Galore v0.6.5 for software cutadapt) and maps reads (software STAR v2.7.3a). Alignments were then post-processed using two workflows to improve their quality: *abra-nf* (v3.0) performs local realignment using software ABRA2 (v2.22^27^), and *BQSR-nf* (v1.1) performs base quality score recalibration using GATK (v4.0.5.1^28^).

Gene- and transcript-level expression quantification was performed using StringTie (v2.1.2^29^) implemented in the workflow *RNAseq-transcript-nf* (v2.2). It generates raw gene count matrices as well as Transcripts Per Million (TPM) and Fragments Per Kilobase Per Million (FPKM) values. Quality control was performed using FastQC (v0.11.9^15^) for raw reads and RSeQC (v3.0.1^30^) for alignment quality.

#### DNA Methylation

IDAT files were processed using R package minfi (v1.40^31^), following our standard workflow Methylation_analysis_scripts. This workflow uses functional normalisation to produce methylation beta- and M-values and incorporates quality control of both samples and probes.

#### Spatial transcriptomics

Samples were processed using Space Ranger (v1.3.0). Data were demultiplexed and reads mapped to the reference genome GRCh38, and tissue and fiducial detection were performed before barcode/unique molecular identifier (UMI) counting, generating feature-barcode matrices. Quality control was performed for both raw and processed data (percentage of valid barcodes and UMIs, quality scores, percentage of reads mapped, median counts per spot).

#### Spatial Proteomics

Data were imported into the GeoMx® Digital Spatial Profiling Data Analysis suite for initial quality control, following the manufacturer’s instructions. AOIs were examined and filtered based on insufficient tissue content, low nuclei count, inadequate segmented area, suboptimal AOI design and poor signal relative to housekeeping or negative control probes. Protein-level quality assessment was conducted using signal-to-background ratios relative to negative control probes.

## Data Records

Processed molecular data (somatic variants, gene expression, methylation M and beta values) are deposited at Zenodo available as *RData* objects or plain-text matrices (csv/tsv). The repository includes somatic variant calls from WGS, bulk RNAseq gene expression matrices, and DNA methylation normalised M- and beta values; for example, objects with *dataset_snv* contain small nucleotide variants, *dataset_purple* copy-number profiles, *dataset_sv* structural variants, *gene_count_matrix* raw RNAseq counts, *gene_TPM_matrix* TPM-normalised matrix, *quanTIseq_matrix* estimated cell type proportions, *Normalised_*Tables* EPIC methylation M- and beta values. Files names may also include samples acronyms (PCA for pulmonary carcinoids, ITH for intra-tumor heterogeneity multi-regional sequencing PCA samples, TR for replicates, LCNEC for large-cell neuroendocrine carcinoma, SCLC for small cell lung cancer). The Zenodo repository also contains the processed data of the four spatial transcriptomics samples from 10X Genomics Visium, provided as scanpy.*rds* objects (*dataset_spatial*) and associated IRIS outputs (*dataset_spatial_IRIS_object*_*20_domains, dataset_spatial_IRIS_annotations, dataset_spatial_IRIS_proportions)*. A cohort level metadata file (*metadata)* is also provided, derived from main study metadata providing information about sample identifiers, histopathological subtype, grade, molecular group labels (Ca A1, Ca A2, Ca B, supra-carcinoid enriched) and other related information. Corresponding raw sequencing data-fastq files for RNA-seq (EGAD00001015524), cram files for WGS (EGAD00001015520), germline variant calls (EGAD00001015672) and idat files for methylation arrays (EGAD00010002480) are hosted on the European Genome-Phenome Archive website, study EGAS00001005979. For these sensitive data that require access controls, authorization for access must be requested to the data access committee (EGAC00001002636) and will require the signing of a Data Transfer Agreement (template is provided in Supplementary information File S1) in order to ensure the respect of the patient’s consent. Additionally, raw spatial transcriptomics data is hosted at the EMBL Spatial transcriptomics portal, and WSI in Aperio *svs* format from the lungNENomics project is hosted in the EBI bioImage Archive (https://doi.org/10.6019/S-BIAD3143).

## Technical validation

### Quality Control

For each data type, we performed quality controls (QC) of raw and processed data. Software MultiQC (v1.31) was used to aggregate the QC results across samples and generate interactive plots. QC plots for WGS (Fig. 1a-c), RNA-seq (Fig. 2a-f,h), and Visium (Fig. 4a-d) were generated by multiQC. MultiQC reports are available in Supplementary Information (files S2-S7) for detailed quality control metrics.

**Fig. 1.**
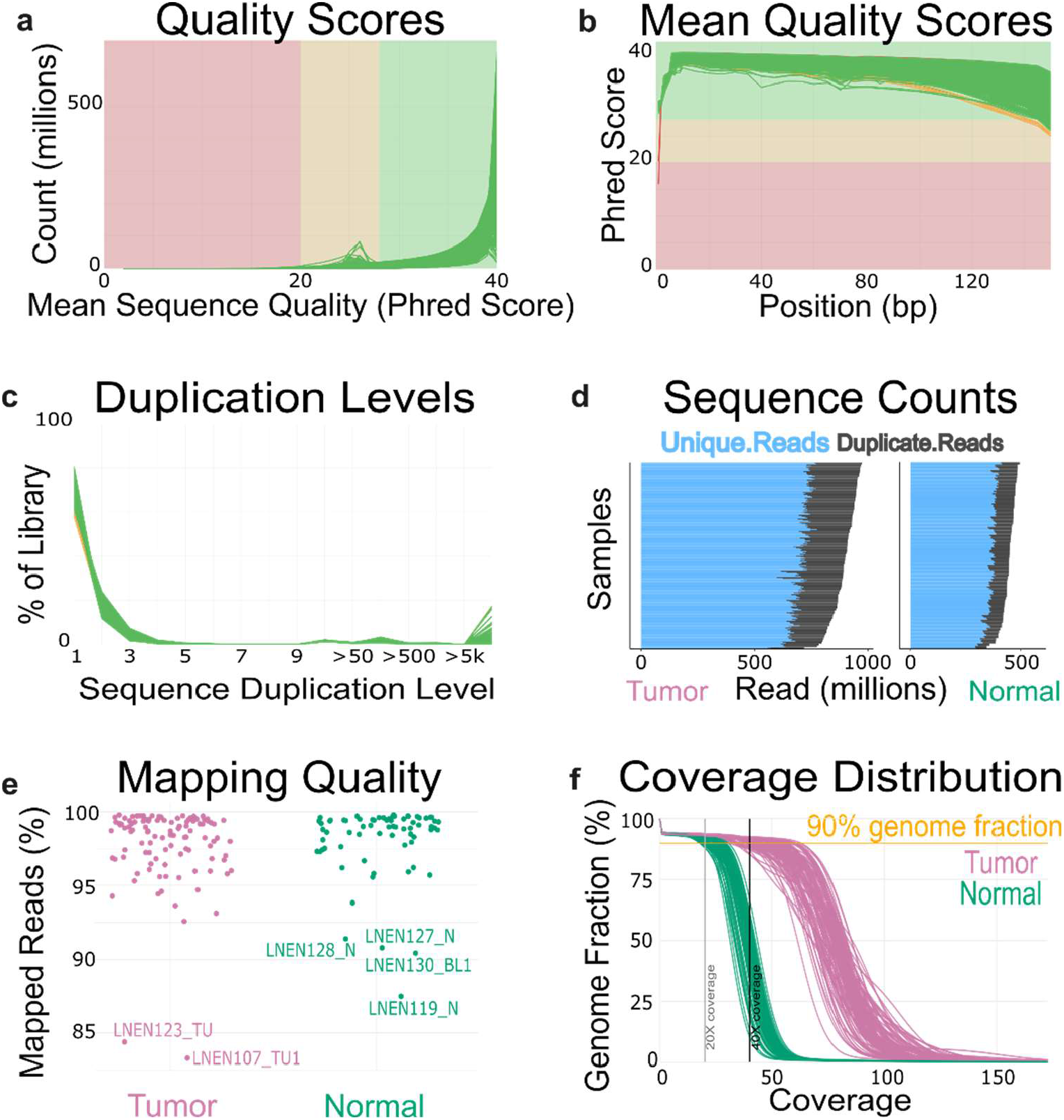
Quality control of the Whole-Genome Sequencing (WGS) data. (a) Distribution of the mean sequence quality of the reads in Phred score. (b) Mean sequence quality score per position in the read in base pairs (bp). (c) Percentage of the library with a given level of duplication. (d) Number of unique and duplicated reads per group (tumour, normal) per file. (e) Percentage of mapped reads per group (tumour and matched normal). (f) Genome fraction as a function of sequencing coverage, with the dashed line indicating 90% genome coverage. In panels (a-c), each line corresponds to a fastq file, with each of the 72 samples (72 tumour, 72 normal, 34 multi-region tumour samples) for a total of 2×2×72 + 2×34 = 356 files; in panel (d), each horizontal bar corresponds to a file. In (a-b), green lines correspond to files that passed the most stringent QC filters of software FastQC; orange lines correspond to files that passed a less stringent filter, and red lines correspond to files that failed the QC filters. In panels (e-f), each point/line corresponds to a sample; pink: tumour, green: normal.

**Fig. 2.**
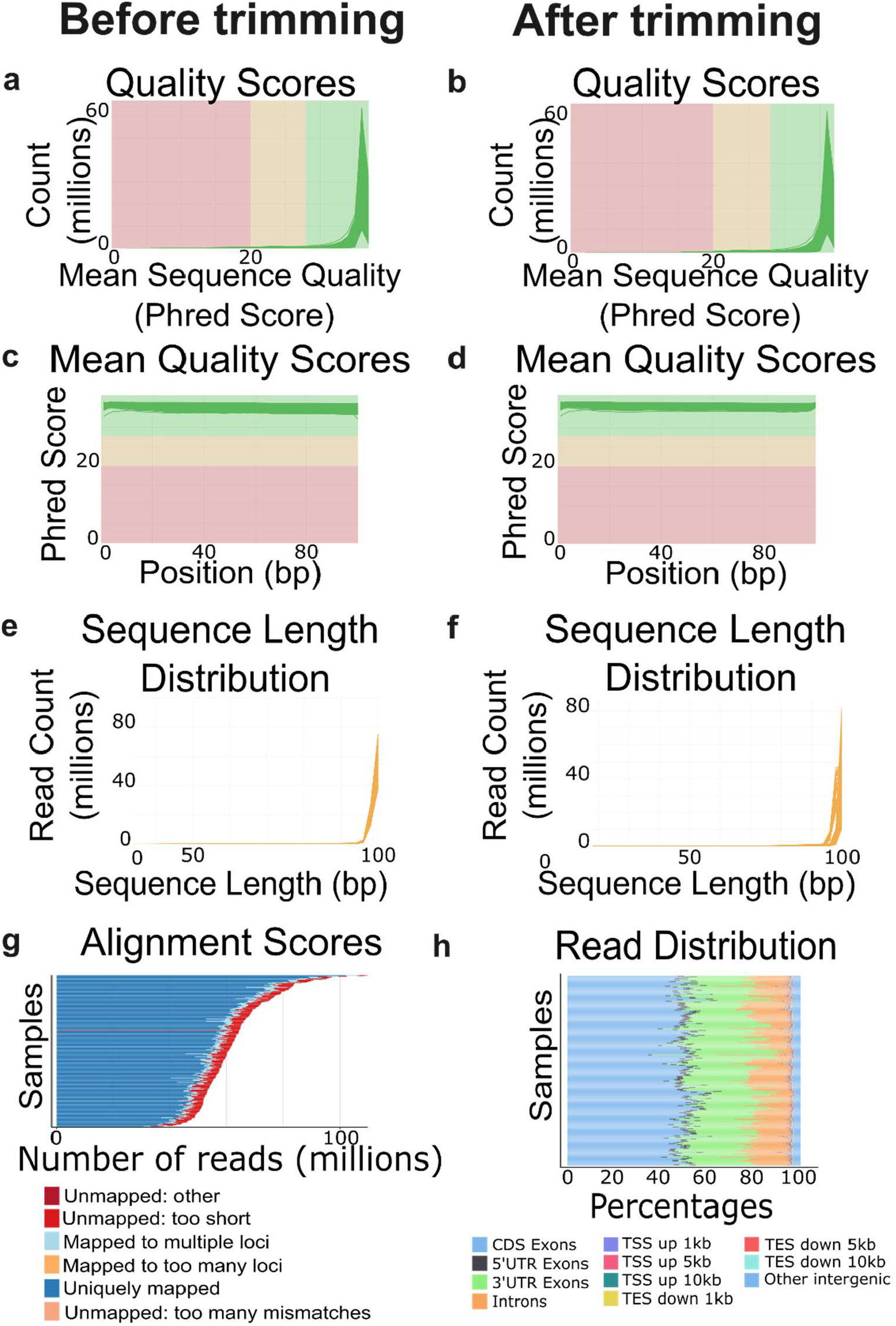
Quality control of the RNA-seq data. Panels (a), (c), (e) correspond to samples before read trimming; panels (b), (d), (f) correspond to controls after read trimming. (a-b) Distribution of mean quality scores of the reads in Phred scale. (c-d) Mean sequence quality score per position in the read in base pairs (bp). (e-f) Sequence read length distributions. (g-h) Quality control of the RNA-seq alignments. (g) Number of sequence tags with each alignment score. (h) Genomic distribution of mapped reads across annotated regions. In panels (a-f), each line corresponds to a fastq file, and was subdivided into two read-pair files, for a total of 2×178 + 2×61 files (multi-region samples), and 2×73 files (technical replicates, corresponding to samples sequenced across multiple lanes or runs), for a total of 624 fastq files. In panels (g-h), each line corresponds to one BAM file (178 + 61 = 239 files).

#### WGS

Raw reads passed quality control filters across all samples. Most of the libraries displayed high sequence quality scores with a mean Phred score above 30 (Fig.1a and b). A gradual decrease was observed towards the 3’ end of the reads, as expected for Illumina sequencing, while remaining within the acceptable quality ranges. Only two samples (LNEN114_N2, LNEN114_T3) showed lower Phred scores (<30), particularly at the beginning of the read (5’ end) (Fig. 1b). Adapter content was very low across all the samples (<2.5% of reads contained adapter sequences), and hence all passed QC. Sequence duplication levels were within acceptable limits (<30% of reads duplicated), indicating good library quality (Fig. 1c). The number of reads was consistent between read pairs and respected target read depths (Fig. 1d): samples with a target depth of 30x (normal) had a lower number of reads (2×450M = 900M reads) than tumour samples (2×900M reads= 1800M reads), which had a target depth of 60x (tumour). The proportion of mapped reads was consistently high in both tumour and matched normal, with most samples showing mapping rates of more than 95% (Fig. 1e). The tumour and matched normal samples showed a mean coverage of greater than 60x and 30x, respectively, consistent with the intended sequencing depths (Fig. 1f).

#### RNA-seq

Raw reads passed quality filters after read trimming. Mean Quality Scores were consistently high across the samples before and after trimming, with Phred scores more than 30 for the majority of bases (Fig. 2a, b). Phred score per position also remained stable, with trimming not altering overall quality distribution (Fig. 2c, d). All the samples had minimal adapter content after trimming (<0.1%). Overall read length remained largely preserved before and after trimming (Fig. 3e, f). The number of reads was consistent with the target of 50 million reads: reads sequenced on a single lane had at least 2×30M reads, and those sequenced on two or more lanes had at least 4×20M reads. Following read alignment, junction saturation analysis revealed that the fraction of known splice junctions (i.e., junctions annotated in the gencode v33 annotation file) rapidly achieved a plateau as the proportion of reads increased, indicating sufficient sequencing depth to capture most annotated splice events (Supplementary Fig. S1a). In contrast, novel junctions increased more gradually with number of reads, and showed weaker saturation (no complete plateau), indicating more abundant unannotated regions and that rarer events may remain undetected (Supplementary Fig. S1b). Overall, alignment scores were good, with the majority of reads uniquely mapped and only a small fraction assigned to multiple mapping or unmapped categories (Fig. 2g). Unmapped reads were primarily due to short reads. Only one sample (LNEN079_TU) showed a very high number of unmapped reads compared to the rest of the samples; we subsequently investigated how this low read count affected downstream analyses (Data Validation). The genomic read distribution of mapped reads showed that the majority aligned to exons (45-55%), 3’UTRs (20-40%), and introns (5-25%) (Fig. 2h).

**Fig. 3.**
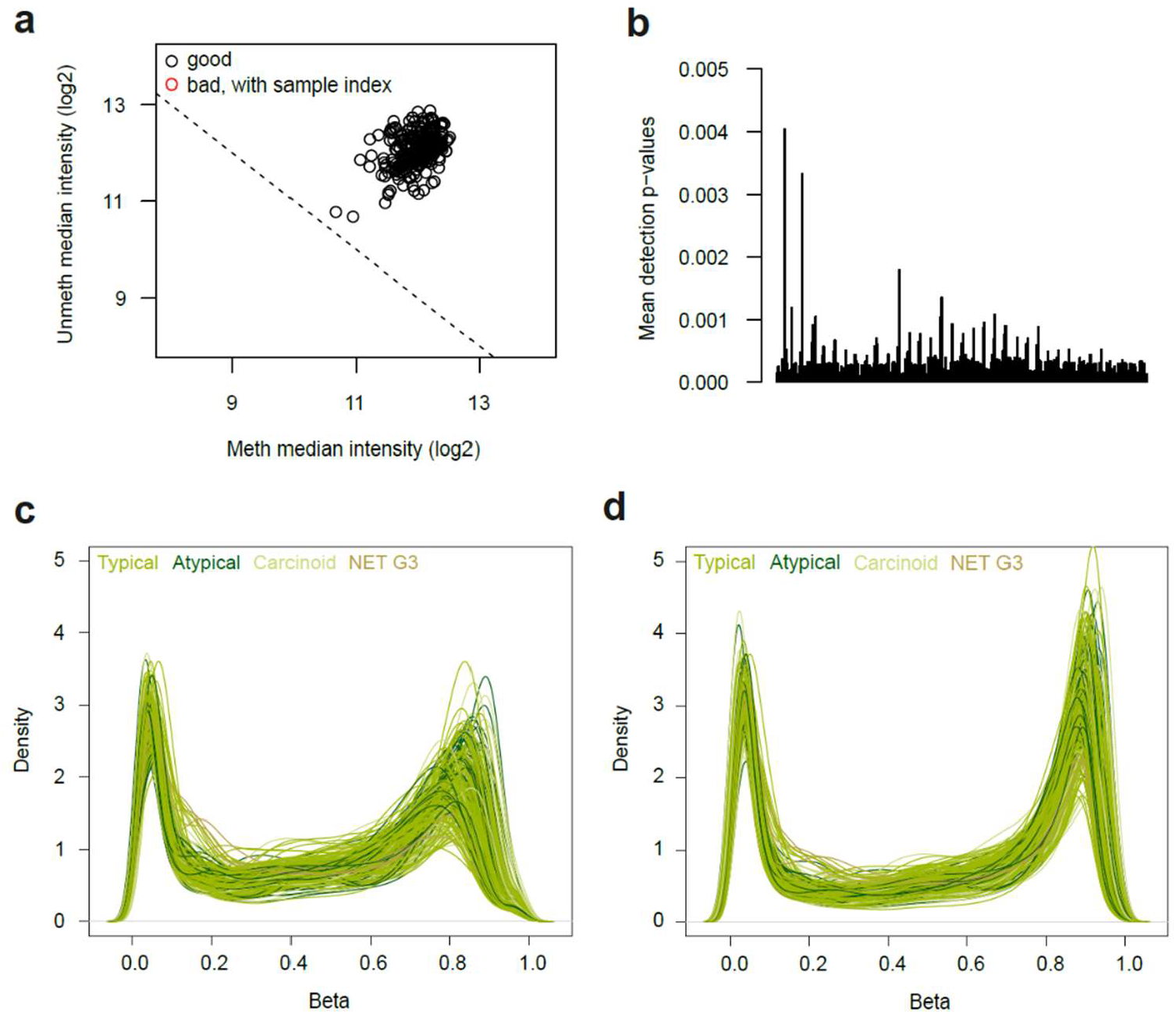
Quality control of DNA methylation data. (a) Log2 methylated (x-axis) and unmethylated (y-axis) signal intensity plot of 277 samples. No samples fell below the cut-off. (b) Mean detection *P* values across probes (y-axis) per sample (x-axis) showing all samples were below the threshold of 0.01 for sample exclusion. (c) Pre-normalisation beta density plot of 277 samples across 865,859 probes, coloured by tumour type, prior to functional normalisation. (d) Beta density plot of 277 samples across 761,463 probes, coloured by tumour type, following functional normalisation and removal of failed probes (*P* detection value > 0.01), cross-reactive probes, sex chromosome probes, single-nucleotide polymorphism probes.

#### DNA Methylation

Two-colour intensity data of internal control probes were manually inspected to check the quality of successive sample preparation steps (bisulphite conversion, hybridisation, extension, and staining; ENmix). All samples passed the QC steps of per sample log2 methylated and unmethylated chip-wise median signal intensity comparison (minfi, Fig. 3a), and overall *P*-detection value measurement (all *P* < 0.01, minfi, Fig. 3b). Following functional normalisation (minfi), quality control was performed on probes. Four groups of probes were removed: (i) poor performing probes with a *P*-detection value > 0.01 in at least one sample (*n* = 38,137), *P*-detection value was computed by comparing the total signal (methylated and unmethylated) of each probe with the background signal level from non-negative control probes (minfi); (ii) cross-reactive probes (*n* = 41,919), cross-reactive probes co-hybridise to multiple locations within the genome and therefore cannot be reliably investigated; (iii) probes on the sex chromosomes (*n* = 16,591), and (iv) probes with SNPs within the single base extension site, or target CpG site, at a minor allele frequency of > 5% (*n* = 7,733, database dbSNP build 137). This resulted in a dataset comprising 761,463 probes for 277 samples (Fig. 3c-d). Beta and M-values were extracted (functions getBeta and getM, minfi), and probes recording M-values of −∞ for at least one sample were replaced with the next lowest M-value in the dataset.

#### Spatial transcriptomics

For raw data, across samples, mean sequence quality was high (>30 Phred score) (Fig. 4a). The majority of base pair (bp) positions were high-quality (>30 Phred score), with a sharp decline in quality from approximately position 50 onward, where Phred scores dropped to between 20 and 30 (Fig. 4b). Median genes detected per spot ranged from 1876 to 6834, with a mean UMI count ranging from 58.3k to 88.4k per spot (Fig. 4c). Each spatial transcriptomics sample quality control of mapped reads was then performed at the spot level. Per-sample thresholds for the metrics of total number of detected UMIs per spot, and the number of detected genes per spot were visualised by red dashed lines on the pre-filter distribution and spatial plots (Fig. 4d-g). The thresholds were determined for each sample from the distribution of each quality control metric where 1% and 99% are determined as outliers and were double-checked manually to avoid removal of tissue-associated spots. Spots that did not meet these criteria were removed from the downstream analysis.

**Fig. 4.**
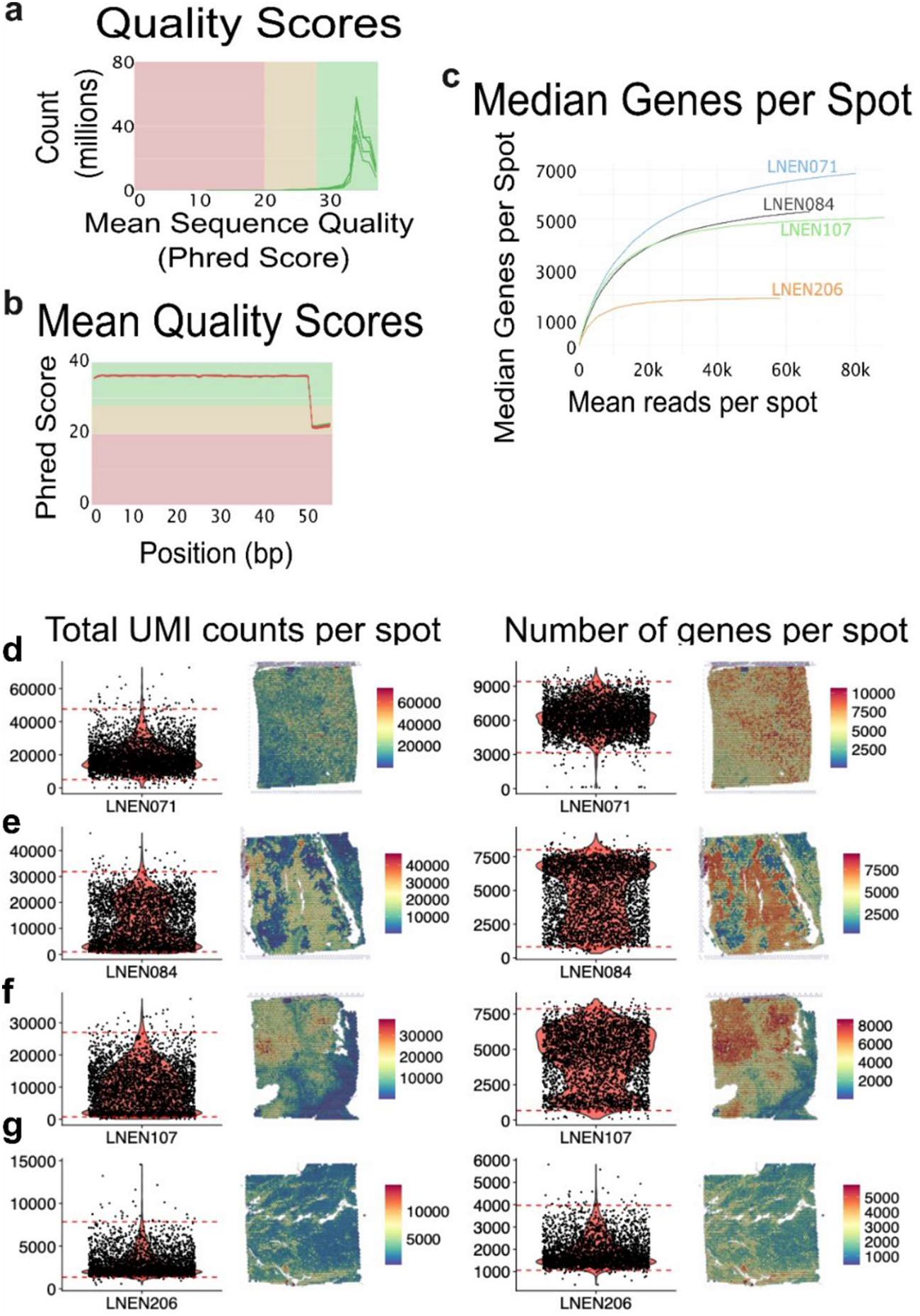
Quality control of the Spatial transcriptomics data. (a) Distribution of mean quality scores of the reads in Phred score. (b) Mean sequence quality score per position in the read in base pairs (bp). Some samples are coloured red because at positions >50, whilst the mean per-base quality remained above 20, the median fell below 20 (the criteria used by FastQC for pass/fail quality assessment). (c) Median gene counts per spot. (d-g) Spot level quality control of Spatial transcriptomics data. Violin plots and spatial maps showing the pre-filter distribution of UMI counts (left panel) and feature counts (right panel) per spot across representative Visium tissue sections; red dashed lines on the violins indicate the per-sample 1% and 99% quantile thresholds; spots outside these bounds were removed prior to downstream analysis; each row corresponds to one representative sample. (d) LNEN071, (e) LNEN084, (f) LNEN107, (g) LNEN206.

#### Spatial Proteomics

Quality control was performed on AOIs and protein probes according to the manufacturer’s instructions (GeoMx® Data Analysis and nCounter user manuals). Of the 513 AOIs examined (Fig. 5a, b), 44 were excluded (8.6%) based on the predefined criteria: mixed tumour/immune region design (*n* = 4), low cellular content (<20 nuclei, *n* = 16), limited AOI surface area (less than 1600µm^2^, *n* = 6), insufficient material collected (*n* = 15), or insufficient expression of housekeeper probes (logarithm of geometric mean less than 5 for S6, Histone H3, GAPDH), or negative control probes (logarithm of geometric mean less than 3 for Rb IgG, Ms IgG2, *n* = 3). This resulted in 469 AOIs for downstream analysis. Protein-level quality assessment was performed by evaluating probe signal relative to background, defined as the geometric mean of negative control probes. Proteins for which the mean log2 signal to background ratio was less than zero were removed. Based on this criteria, nine proteins were excluded from tumour cell AOIs (CD66b, CD163, CD80, PD-L1, CD27, ICOS, PD-L2, CD40, and PD-1) (Fig. 5d), and five from immune cell AOIs (CD80, PD-L2, CD66b, PD-L1, and FOXP3) (Fig. 5e). Only proteins that passed QC in both tumour and immune cell AOIs were retained, leaving 23 proteins for analysis.

**Fig. 5.**
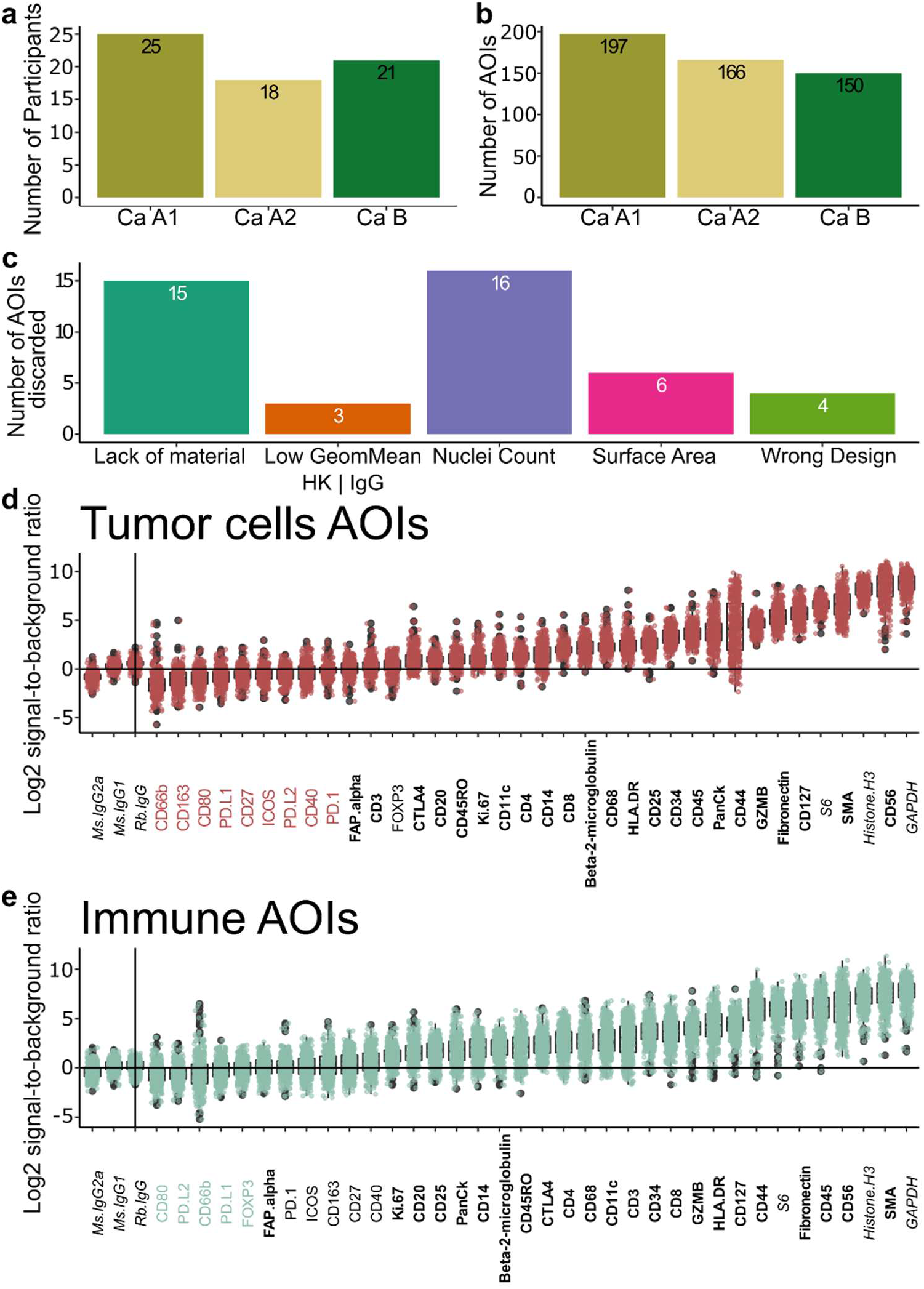
Data description and quality control of the Digital Spatial Profiler (DSP) experiment. (a) Number of patients included in the DSP experiment per molecular group. (b) Number of areas of interest (AOIs) per molecular group selected for the DSP experiment. (c) Quality control of the AOIs and reasons for exclusion of 44 AOIs. (d) Log2 distributions of the signal to background ratio, within the tumour cells AOIs. The background was estimated as the geometric mean of the negative control probes. The signal from the IgG probes is indicated to the left of the vertical bar. The name of the protein in colour corresponds to the protein that was excluded from the analysis according to the following criteria: mean log2 signal/background ratio less than 0. (e) As per (d) for immune cells AOIs.

**Fig. 6.**
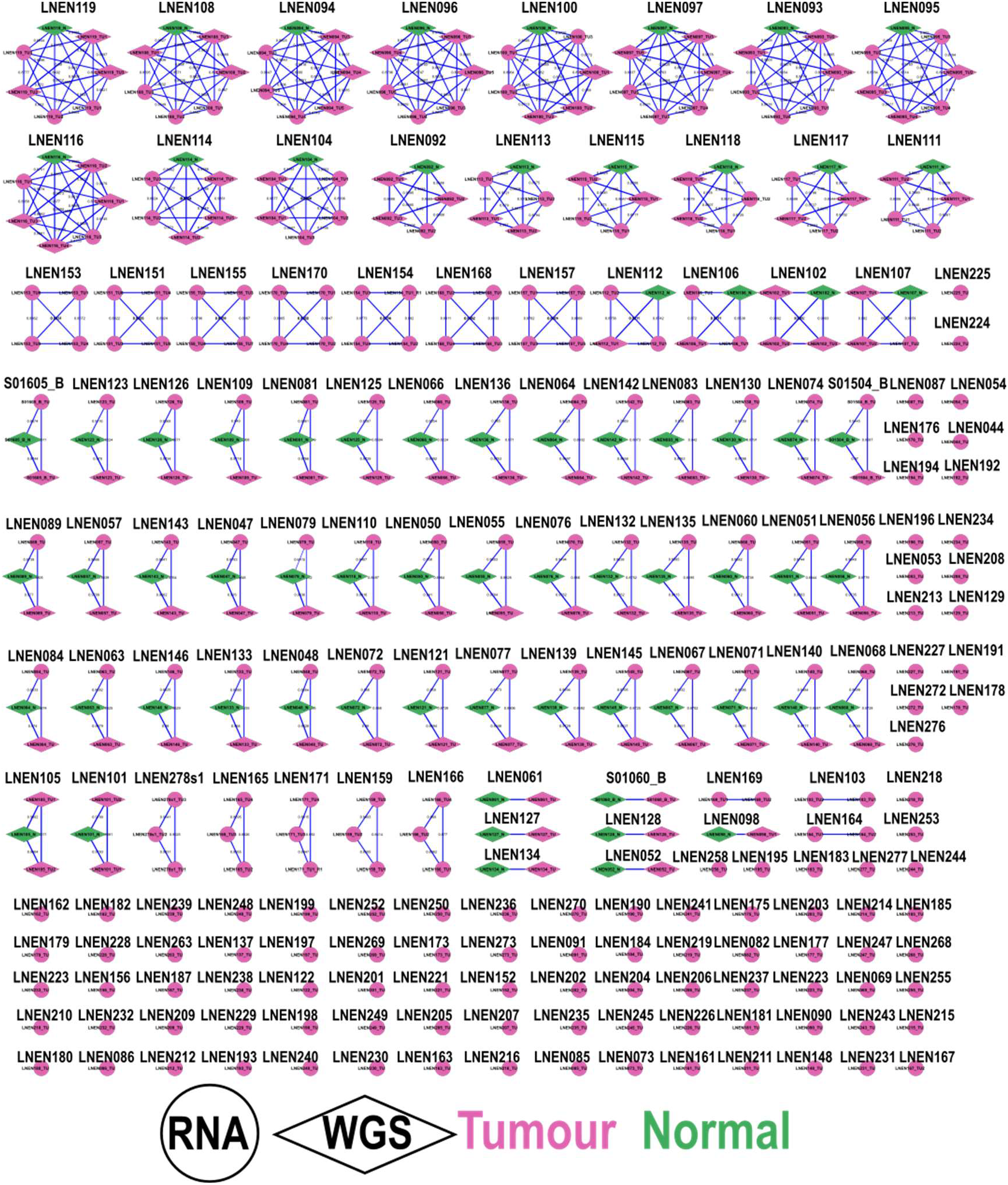
Network of matches between tumour, ITH and matched normal samples of WGS, computed with NGSCheckMate. Numbers on the edges and edge thickness correspond to the Pearson correlation coefficient (*r*) between allelic fractions for the germline SNP panel. Shapes indicate experiments; circle: RNAseq, diamond: WGS; colour indicates condition; pink: tumour, green: normal

### Data Validation

*Sample matching*. We used software NGSCheckMate^32^ to check that samples from the same experiment indeed came from the same individual in WGS and RNAseq, using our workflow NGSCheckMate-nf (v1.1). The sample matching algorithm correctly identified all samples from the same individual, based on a correlation between VAF of common germline variants exceeding 0.5 (Fig.6).

#### RNA-seq of LNEN079

LNEN079_TU had lower sequencing depth (~1 million) compared to the rest of the cohort (~50 million) for RNAseq. Because the main goal of the lungNENomics study was to perform a molecular classification of the tumors, we assessed whether molecular classification results were affected by this low sequencing depth. To do so, we selected 10 samples closest to LNEN079_TU in terms of molecular classification as a proxy validation set. The molecular classification was performed using archetype analysis, a type of fuzzy clustering where a number of extreme profiles are first identified (*K*=4 in the lungNENomics study), and each sample is then assigned a proportion for each archetype - a value between 0 and 1 reflecting its closeness to this extreme profile, where the sum of all scores is equal to 1^10^. These archetype proportions thus represent a simple quantitative measurement of the profile of each tumor. To find the 10 samples closest to LNEN079_TU, we computed the Euclidean distance between all samples in terms of archetype proportions and selected the samples with the shortest distance (LNEN213_TU, S02339, LNEN184_TU, LNEN74_TU, LNEN187_TU, LNET19T, LNEN030_TU, LNEN151_TU3, LNEN154_TU3, LNENE154_TU3, LNEN211_TU). LNEN030_TU was excluded due to lack of expression data, reducing the proxy set to 9. For each sample, 100 expression vectors were simulated by downsampling raw read counts to LNEN079_TU library size using *rmultinom* in R, thereby mimicking the low-depth sequencing. Each simulated expression vector was reintegrated into the multi-omics pipeline with all other inputs and parameters held constant, and the resulting MOFA factors values and archetype proportions were compared to those derived from original full depth data.

MOFA2 factor values and archetype proportions were robustly recovered across 100 simulated low-depth iterations for each of the 9 validation samples, with high correlation values for MOFA latent factors (Factor1, Factor2, Factor5; *r* >0.9, Supplementary figure S2) and RMSE values consistently below 0.1 across all four archetypes (sc-enriched, Ca A1, Ca A2, Ca B; Supplementary figure S3) demonstrating that MOFA-based projection and archetype assignment of all downsampled samples are stable across all iterations (Supplementary figure S4).

#### Sex validation

We validated participant sex reported in clinical data using WGS, RNA-seq, and DNA methylation data. Sample sex was inferred from WGS using the sex determination module implemented in purple, based on independent assessment from amber (v3.5) and cobalt (v1.11). In cases of discordance between both the tools, cobalt-based assignment was retained and flagged as failed sex validation. No samples were discordant between clinical and WGS-predicted sex (Fig. 7a). For RNA-seq, the sum of variance stabilised read counts for sex chromosomes were plotted for each tumour sample resulting in two clusters (Fig. 7b). One sample was identified as an outlier, LNEN199_TU, which had very low chromosome Y expression for its clinical sex (male). For methylation data, participant sex was predicted based on median total intensity of sex chromosome signal (R package minfi). One sample was discordant between clinical and DNA methylation predicted sex, LNEN199_TU, having lower chromosome Y intensity than expected for a male sample (Fig. 7c). To further investigate the discordant sample, its chromosome Y expression and DNA methylation data were compared (Fig. 7d), finding that LNEN199_TU is likely female given the low chromosome Y values in both data types. Clinical data for LNEN199_TU were likely entered erroneously and were therefore excluded from further analysis.

**Fig. 7.**
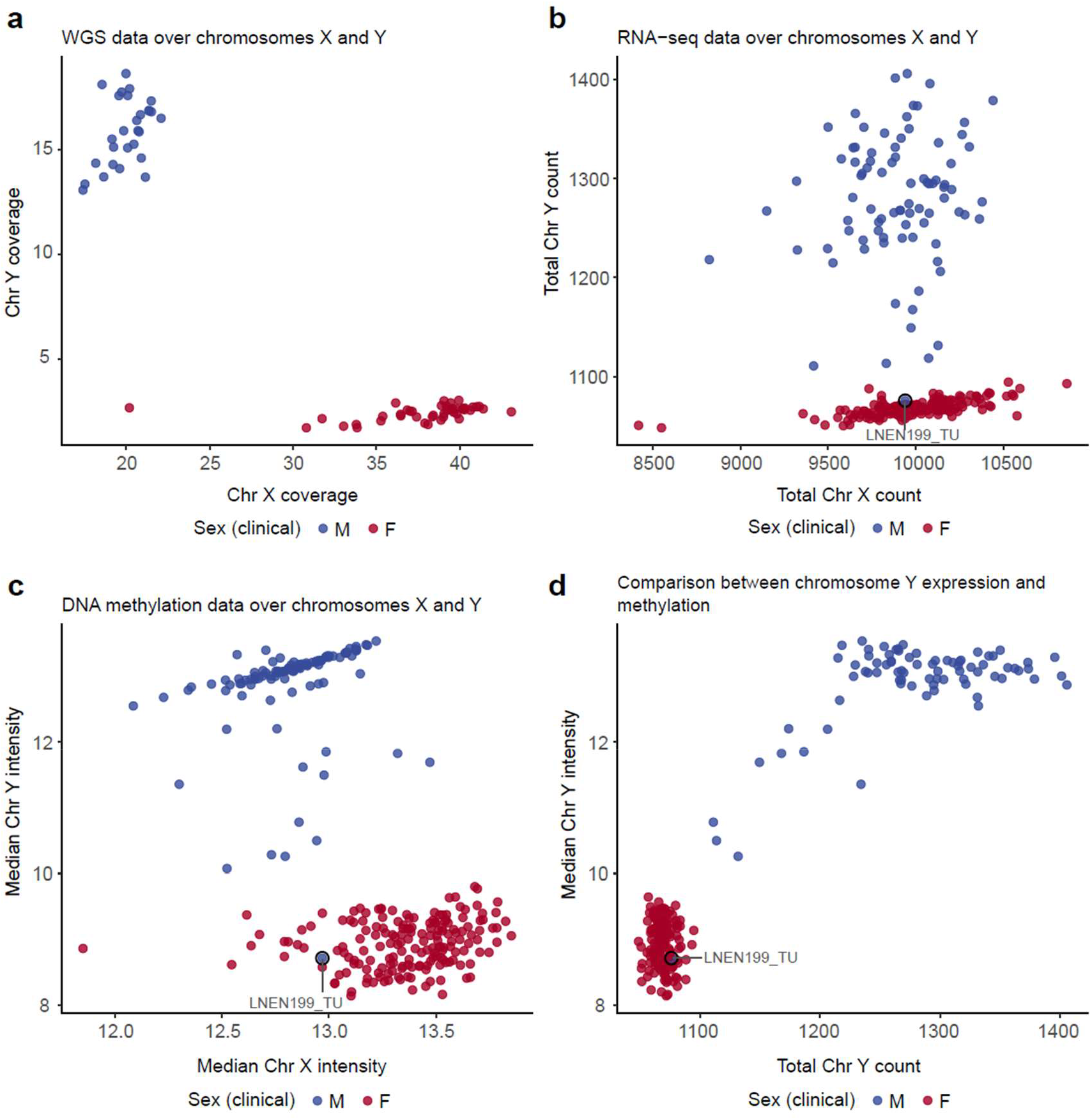
Sex validation. (a) Chromosome X (x-axis) and Y (y-axis) coverage from WGS data per matched normal sample (*n* = 72). (b) Total expression level of the X (x-axis) and Y (y-axis) chromosomes (sum of variance-stabilised read counts) per sample (*n* = 239). (c) Median DNA methylation array total intensity on the X (x-axis) and Y (y-axis) chromosomes per sample (*n* = 277). (d) Comparison between total Y chromosome expression (x-axis) and median DNA methylation array total Y chromosome intensity (y-axis) for each sample profiled by both RNA sequencing and DNA methylation (*n* = 225). Point colours correspond to clinically reported sex, M, male; F, female. Samples circled and labelled in black were discordant between clinically reported sex and predicted sex by at least one omic type.

## Data Availability

Processed molecular data are openly available at Zenodo under a Creative Commons Attribution 4.0 International license. Raw molecular data - WGS cram files (EGAD00001015520), RNAseq fastq files (EGAD00001015524), germline variant calls (EGAD00001015672), and idat files for DNA methylation (EGAD00010002480) are deposited under control access European Genome-Phenome Archive website, study EGAS00001005979. Access to this data must be requested from the Data Access Committee (https://ega-archive.org/dacs/EGAC00001001811) and requires a completion of a Data Transfer Agreement (template is provided in Supplementary information File S1) to ensure respect with patient consent and data protection regulations. Raw spatial transcriptomics data is hosted at the EMBL Spatial transcriptomics portal and medical imaging data (Hematoxylin & Eosin stained whole-slide images) from the project is hosted in the EBI bioImage Archive (10.6019/S-BIAD3143).

## Code Availability

All pipelines used in this study are openly accessible at IARCbioinfo. Default parameters from the original developers were applied when not explicitly specified in the Methods. All custom scripts and reproducible code supporting the analyses are available at https://github.com/IARCbioinfo/MS_lungNENomics.

## Author Contributions

Conceptualization: L.F-C., M.F., N.A.

Data curation: L.K., A.S-O., E.M., C.V., A.DG., Z.L., N.A.

Funding acquisition: L.M., L.F-C., M.F., N.A.

Methodology: E.M., N.A.

Project administration: A.S-O., L.F-C., M.F.

Software: L.K., A.S-O., E.M., A.DG., Z.L., N.A.

Supervision: J.K., L.M., L.B., L.F-C., M.F., N.A.

Validation: L.K., A.S-O., E.M., N.A.

Visualization: L.K., A.S-O., E.M., Z.L., N.A.

Writing – original draft: L.K., A.S-O., E.M., L.F-C., M.F., N.A.

Writing – review & editing: all authors

## Acknowledgements

The lungNENomics project is part of the Rare Cancers Genomics initiative (www.rarecancersgenomics.com/) led by the Computational Cancer Genomics team at the IARC (https://www.iarc.who.int/teams-ccg/). We thank the Hospices Civils de Lyon (CRB-HCL BB-0033-00046) and Centre Léon Bérard (CRB-CLB BB-0033-00050) biobanks in Lyon, as well as Côte d’Azur University’s biobank (BB-0033-00025) in Nice, France, all authorised by the French Ministry of Research, for sharing human biological samples and associated data. We thank the 12 centres that voluntarily participated in the lungNENomics project by providing one FFPE block per patient: The Tumour Bank of the François Baclesse Centre in Caen (France), the Institut für Diagnostik und Forschung in Pathologie of the Medical University of Graz (Austria), the Department of Biopathology of the Léon Bérard Centre in Lyon (France), the Institute of Pathology of the Hospices civils de Lyon (France), the Department of Surgical Oncology of St Vincent’s Hospital in Melbourne (Australia), the Department of Oncology and Haemato-Oncology of the University of Milan (Italy), the Department of Biopathology at the Nancy Regional Hospital (France), the Laboratory of Clinical and Experimental Pathology at the Pasteur Hospital in Nice (France), the Departments of Pathology and Oncology at Oslo University Hospital (Norway), the Pathology Department at the Cochin Hospital in Paris (France), the Oncology Unit at the IRCCS Casa Sollievo della Sofferenza Foundation in Rotondo (Italy), and the Oncology Department at the University of Turin (Italy).

## Ethics Statement

This study was approved by the International Agency for Research on Cancer Ethics Committee (project number 19-07).

