## Supplementary Information (files S2-S7) for "A multi-omic, spatial, and whole-slide image dataset of lung neuroendocrine tumours from the lungNENomics cohort": S4.html

LUNGNENOMICS: MultiQC Report


### Toggle navigation v1.31

### LUNGNENOMICS

Loading report..

- General Stats
- FastQC
  - Sequence Counts
  - Sequence Quality Histograms
  - Per Sequence Quality Scores
  - Per Base Sequence Content
  - Per Sequence GC Content
  - Per Base N Content
  - Sequence Length Distribution
  - Sequence Duplication Levels
  - Overrepresented sequences by sample
  - Top overrepresented sequences
  - Adapter Content
  - Status Checks
- Software Versions

Toolbox

##### MultiQC Toolbox

###### Apply Highlight Samples

+

Regex mode off
help
 Clear

###### Apply Rename Samples

+

Click here for bulk input.

Paste two columns of a tab-delimited table here (eg. from Excel).

First column should be the old name, second column the new name.

Add

Regex mode off
help
 Clear

###### Apply Show / Hide Samples

Hide matching samples

Show only matching samples

+

Regex mode off
help
 Clear

###### Explain with AI

Configure AI settings to get explanations of plots and data in this report.

AI Provider

Endpoint


Use the OpenAI API-style requests with a custom endpoint.

Model

API Key


Keys entered here will be stored in your browser's local storage.
See the docs.

Additional Payload


Any additional options passed in API request payload. Enter as a JSON object.

Context Window


The maximum number of tokens that can be processed in a single request


---

Anonymize samples off

###### Export Plots

- Images
- Data

px

px

Aspect ratio

PNG
SVG

Plot scaling

X

Download the raw data used to create the plots in this report below:

Format:

Tab-separated
Comma-separated
JSON

Note that additional data was saved in `multiqc_data` when this report was generated.

---

###### Choose Plots

 All
 None


---


   Download Plot Images

If you use plots from MultiQC in a publication or presentation, please cite:

Loading report..

Report
generated on 2025-12-08, 14:44 CET
based on data in:
`/data/lungNENomics/work/lipikal/Gigascience/figures_061225/RNAseq/selected_samples/pretrim`

Summarize report

Copy report prompt

Change sample names:
Sequencing Center ID
lungNENomics\_ID

---

×
don't show again

**Welcome!** Not sure where to start?  
Watch a tutorial video
  *(6:06)*

×

Because this report contains a lot of samples, you may need to click 'Show plot' to see some graphs.
Render all plots

**Report AI Summary**

More details...

Provider: ,
model: 

Chat with Seqera AI

#### General Statistics

**AI Summary**

Provider: ,
model:

Chat with Seqera AI

Showing violin plots for 624 data points.

Export...

Copy prompt


Summarize plot

Created with MultiQC

×

###### General Statistics: Columns

**AI Summary**

Provider: ,
model:

Chat with Seqera AI

Export...

Copy prompt


Summarize plot

Created with MultiQC

---

##### Sequence Length Distribution

The distribution of fragment sizes (read lengths) found. See the FastQC help

**AI Summary**

Copy Prompt


Summarize table

| Overrepresented sequence | Reports | Occurrences | % of all reads |
| --- | --- | --- | --- |
| GGGGGGGGGGGGGGGGGGGGGGGGGGGGGGGGGGGGGGGGGGGGGGGGGG | 307 | 67448139 | 0.2223% |
| GTCAGGAGTTCTCAGCTTTCACCAAAAGGTCAGAAGTTATTGCAGTTGTG | 54 | 4598129 | 0.0152% |
| CCCGTATCGAAGGCCTTTTTGGACAGGTGGTGTGTGGTGGCCTTGGTATG | 27 | 1749843 | 0.0058% |
| GGTGTATGCATCGGGGTAGTCCGAGTAACGTCGGGGCATTCCGGATAGGC | 22 | 1317031 | 0.0043% |
| CACATGCCTATCATATAGTAAAACCCAGCCCATGACCCCTAACAGGGGCC | 19 | 1290977 | 0.0043% |
| CCTAGACCAAACCTACGCCAAAATCCATTTCACTATCATATTCATCGGCG | 19 | 1102173 | 0.0036% |
| GTCAGAAGTTATTGCAGTTGTGCCCAGTGGATAGGATGGAGGAAGGGAAA | 23 | 1006127 | 0.0033% |
| GTTAGAGAAATGAATGAGCCTACAGATGATAGGATGTTTCATGTGGTGTA | 17 | 959831 | 0.0032% |
| ATCACATGCCTATCATATAGTAAAACCCAGCCCATGACCCCTAACAGGGG | 12 | 858280 | 0.0028% |
| CCTCAGAGTACTTCGAGTCTCCCTTCACCATTTCCGACGGCATCTACGGC | 14 | 820040 | 0.0027% |
| GTTTGGATGTAAAGTGAAATATTAGTTGGCGGATGAAGCAGATAGTGAGG | 12 | 801057 | 0.0026% |
| GTTGGTTAGTAGGCCTAGTATGAGGAGCGTTATGGAGTGGAAGTGAAATC | 12 | 779567 | 0.0026% |
| GTTTAGGAGTGGGACTTCTAGGGGATTTAGCGGGGTGATGCCTGTTGGGG | 14 | 698076 | 0.0023% |
| GTCAGGAGTAGGAGACAAGGCGCGTAGGGGGCGATAGGGTGGCCTGGGCT | 19 | 696392 | 0.0023% |
| GTGGAAGTGAGCTACAACGTAGTACGTGTCGTGTAGTACGATGTCTAGTG | 12 | 631435 | 0.0021% |
| AGAGAAATGAATGAGCCTACAGATGATAGGATGTTTCATGTGGTGTATGC | 13 | 622719 | 0.0021% |
| GTCGGAAATGGTGAAGGGAGACTCGAAGTACTCTGAGGCTTGTAGGAGGG | 12 | 611182 | 0.0020% |
| CAGGAGTTCTCAGCTTTCACCAAAAGGTCAGAAGTTATTGCAGTTGTGCC | 14 | 548203 | 0.0018% |
| GTCAATGTTTAAAGTTGTGGGGCTGCCCTGCAAAGGATGTTCCAGGGCAG | 14 | 475462 | 0.0016% |
| GTTCAGAGAAGGAATCGTCAATGTTTAAAGTTGTGGGGCTGCCCTGCAAA | 15 | 456233 | 0.0015% |

Close
