## Supplementary Information (files S2-S7) for "A multi-omic, spatial, and whole-slide image dataset of lung neuroendocrine tumours from the lungNENomics cohort": S5.html

LUNGNENOMICS: MultiQC Report


### Toggle navigation v1.31

### LUNGNENOMICS

Loading report..

- General Stats
- RSeQC
  - Read Distribution
  - Junction Saturation
- Cutadapt
  - Filtered Reads
  - Trimmed Sequence Lengths (3')
- FastQC
  - Sequence Counts
  - Sequence Quality Histograms
  - Per Sequence Quality Scores
  - Per Base Sequence Content
  - Per Sequence GC Content
  - Per Base N Content
  - Sequence Length Distribution
  - Sequence Duplication Levels
  - Overrepresented sequences by sample
  - Top overrepresented sequences
  - Adapter Content
  - Status Checks
- Software Versions

Toolbox

##### MultiQC Toolbox

###### Apply Highlight Samples

+

Regex mode off
help
 Clear

###### Apply Rename Samples

+

Click here for bulk input.

Paste two columns of a tab-delimited table here (eg. from Excel).

First column should be the old name, second column the new name.

Add

Regex mode off
help
 Clear

###### Apply Show / Hide Samples

Hide matching samples

Show only matching samples

+

Regex mode off
help
 Clear

###### Explain with AI

Configure AI settings to get explanations of plots and data in this report.

AI Provider

Endpoint


Use the OpenAI API-style requests with a custom endpoint.

Model

API Key


Keys entered here will be stored in your browser's local storage.
See the docs.

Additional Payload


Any additional options passed in API request payload. Enter as a JSON object.

Context Window


The maximum number of tokens that can be processed in a single request


---

Anonymize samples off

###### Export Plots

- Images
- Data

px

px

Aspect ratio

PNG
SVG

Plot scaling

X

Download the raw data used to create the plots in this report below:

Format:

Tab-separated
Comma-separated
JSON

Note that additional data was saved in `multiqc_data` when this report was generated.

---

###### Choose Plots

 All
 None


---


   Download Plot Images

If you use plots from MultiQC in a publication or presentation, please cite:

Loading report..

Report
generated on 2025-12-08, 14:14 CET
based on data in:
`/data/lungNENomics/work/lipikal/Gigascience/figures_061225/RNAseq/selected_samples/posttrim`

Summarize report

Copy report prompt

Change sample names:
Sequencing Center ID
lungNENomics\_ID

---

×
don't show again

**Welcome!** Not sure where to start?  
Watch a tutorial video
  *(6:06)*

×

Because this report contains a lot of samples, you may need to click 'Show plot' to see some graphs.
Render all plots

**Report AI Summary**

More details...

Provider: ,
model: 

Chat with Seqera AI

#### General Statistics

**AI Summary**

Provider: ,
model:

Chat with Seqera AI

Showing violin plots for 1248 data points.

Export...

Copy prompt


Summarize plot

Created with MultiQC

×

###### General Statistics: Columns

Uncheck the tick box to hide columns. Click and drag the handle on the left to change order. Table ID: `general_stats_table_table`

Show All
Show None

| Sort | Visible | Group | Column | Description | ID | Scale |
| --- | --- | --- | --- | --- | --- | --- |
| \|\| |  | Cutadapt | Trimmed bases | % total base pairs trimmed | `cutadapt-percent_trimmed` |  |
| \|\| |  | FastQC | Dups | % duplicate reads | `fastqc-percent_duplicates` |  |
| \|\| |  | FastQC | GC | Average % GC content | `fastqc-percent_gc` |  |
| \|\| |  | FastQC | Avg len | Average read length | `fastqc-avg_sequence_length` |  |
| \|\| |  | FastQC | Median len | Median read length | `fastqc-median_sequence_length` |  |
| \|\| |  | FastQC | Failed | Percentage of modules failed in FastQC report (includes those not plotted here) | `fastqc-percent_fails` |  |
| \|\| |  | FastQC | Seqs | Total sequences (millions) | `fastqc-total_sequences` | read\_count |

Close

#### RSeQC

Evaluates high throughput RNA-seq data.*URL: http://rseqc.sourceforge.net**DOI: 10.1093/bioinformatics/bts356*

##### Read Distribution

Read Distribution calculates how mapped reads are distributed over genome features.

**AI Summary**

Provider: ,
model:

Chat with Seqera AI

Percentages
 Export...

Copy prompt


Summarize plot

Created with MultiQC

---

##### Junction Saturation

Junction Saturation
counts the number of known splicing junctions that are observed
in each dataset. If sequencing depth is sufficient, all (annotated) splice junctions should
be rediscovered, resulting in a curve that reaches a plateau. Missing low abundance splice
junctions can affect downstream analysis.

**AI Summary**

Provider: ,
model:

Chat with Seqera AI

All Junctions
Known Junctions
Novel Junctions

 Export...

Copy prompt


Summarize plot

Created with MultiQC

---

#### Cutadapt

*Version:* 
 `2.10`

Finds and removes adapter sequences, primers, poly-A tails, and other types of unwanted sequences.*URL: https://cutadapt.readthedocs.io**DOI: 10.14806/ej.17.1.200*

##### Filtered Reads

This plot shows the number of reads (SE) / pairs (PE) removed by Cutadapt.

**AI Summary**

Provider: ,
model:

Chat with Seqera AI

Percentages
 Export...

Copy prompt


Summarize plot

Created with MultiQC

---

##### Trimmed Sequence Lengths (3') Help

This plot shows the number of reads with certain lengths of adapter trimmed for the 3' end.

Obs/Exp shows the raw counts divided by the number expected due to sequencing errors.
A defined peak may be related to adapter length.

See the cutadapt documentation
for more information on how these numbers are generated.

**AI Summary**

Provider: ,
model:

Chat with Seqera AI

Counts
Obs/Exp

 Export...

Copy prompt


Summarize plot

Created with MultiQC

---

#### FastQC

Copy Prompt


Summarize table

| Overrepresented sequence | Reports | Occurrences | % of all reads |
| --- | --- | --- | --- |
| GGGGGGGGGGGGGGGGGGGGGGGGGGGGGGGGGGGGGGGGGGGGGGGGGG | 267 | 40651937 | 0.1350% |
| GTCAGGAGTTCTCAGCTTTCACCAAAAGGTCAGAAGTTATTGCAGTTGTG | 53 | 4512340 | 0.0150% |
| CCCGTATCGAAGGCCTTTTTGGACAGGTGGTGTGTGGTGGCCTTGGTATG | 27 | 1735535 | 0.0058% |
| GGTGTATGCATCGGGGTAGTCCGAGTAACGTCGGGGCATTCCGGATAGGC | 22 | 1302141 | 0.0043% |
| GGGGGGGGGGGGGGGGGGGGGGGGGGG | 17 | 1300937 | 0.0043% |
| CACATGCCTATCATATAGTAAAACCCAGCCCATGACCCCTAACAGGGGCC | 19 | 1287259 | 0.0043% |
| CCTAGACCAAACCTACGCCAAAATCCATTTCACTATCATATTCATCGGCG | 19 | 1095059 | 0.0036% |
| GTCAGAAGTTATTGCAGTTGTGCCCAGTGGATAGGATGGAGGAAGGGAAA | 23 | 992470 | 0.0033% |
| GGGGGGGGGGGGGGGGGGGGGGGGGG | 13 | 979873 | 0.0033% |
| GTTAGAGAAATGAATGAGCCTACAGATGATAGGATGTTTCATGTGGTGTA | 17 | 948104 | 0.0031% |
| CCTCAGAGTACTTCGAGTCTCCCTTCACCATTTCCGACGGCATCTACGGC | 15 | 877748 | 0.0029% |
| GTTTGGATGTAAAGTGAAATATTAGTTGGCGGATGAAGCAGATAGTGAGG | 12 | 791592 | 0.0026% |
| GGGGGGGGGGGGGGGGGGGGGGGGG | 13 | 776929 | 0.0026% |
| GTTGGTTAGTAGGCCTAGTATGAGGAGCGTTATGGAGTGGAAGTGAAATC | 12 | 776127 | 0.0026% |
| GTTTAGGAGTGGGACTTCTAGGGGATTTAGCGGGGTGATGCCTGTTGGGG | 14 | 688095 | 0.0023% |
| GTCAGGAGTAGGAGACAAGGCGCGTAGGGGGCGATAGGGTGGCCTGGGCT | 19 | 687561 | 0.0023% |
| AGAGAAATGAATGAGCCTACAGATGATAGGATGTTTCATGTGGTGTATGC | 13 | 613044 | 0.0020% |
| CAGGAGTTCTCAGCTTTCACCAAAAGGTCAGAAGTTATTGCAGTTGTGCC | 14 | 540605 | 0.0018% |
| GTCAATGTTTAAAGTTGTGGGGCTGCCCTGCAAAGGATGTTCCAGGGCAG | 14 | 471144 | 0.0016% |
| GTTCAGAGAAGGAATCGTCAATGTTTAAAGTTGTGGGGCTGCCCTGCAAA | 15 | 451237 | 0.0015% |

×

###### FastQC: Top overrepresented sequences: Columns

Uncheck the tick box to hide columns. Click and drag the handle on the left to change order. Table ID: `fastqc_top_overrepresented_sequences_table_table`

Show All
Show None

| Sort | Visible | Group | Column | Description | ID | Scale |
| --- | --- | --- | --- | --- | --- | --- |
| \|\| |  |  | Reports | Number of FastQC reports where this sequence is founds as overrepresented | `fastqc-samples` |  |
| \|\| |  |  | Occurrences | Total number of occurrences of the sequence (among the samples where the sequence is overrepresented) | `fastqc-total_count` |  |
| \|\| |  |  | % of all reads | Total number of occurrences as the percentage of all reads (among samples where the sequence is overrepresented) | `fastqc-total_percent` |  |

Close

---

##### Adapter Content Help

The cumulative percentage count of the proportion of your
library which has seen each of the adapter sequences at each position.

Note that only samples with ≥ 0.1% adapter contamination are shown.

There may be several lines per sample, as one is shown for each adapter
detected in the file.

From the FastQC Help:

*The plot shows a cumulative percentage count of the proportion
of your library which has seen each of the adapter sequences at each position.
Once a sequence has been seen in a read it is counted as being present
right through to the end of the read so the percentages you see will only
increase as the read length goes on.*

**AI Summary**

Provider: ,
model:

Chat with Seqera AI

No samples found with any adapter contamination > 0.1%

---

##### Status Checks Help

Status for each FastQC section showing whether results seem entirely normal (green),
slightly abnormal (orange) or very unusual (red).

FastQC assigns a status for each section of the report.
These give a quick evaluation of whether the results of the analysis seem
entirely normal (green), slightly abnormal (orange) or very unusual (red).

It is important to stress that although the analysis results appear to give a pass/fail result,
these evaluations must be taken in the context of what you expect from your library.
A 'normal' sample as far as FastQC is concerned is random and diverse.
Some experiments may be expected to produce libraries which are biased in particular ways.
You should treat the summary evaluations therefore as pointers to where you should concentrate
your attention and understand why your library may not look random and diverse.

Specific guidance on how to interpret the output of each module can be found in the relevant
report section, or in the FastQC help.

In this heatmap, we summarise all of these into a single heatmap for a quick overview.
Note that not all FastQC sections have plots in MultiQC reports, but all status checks
are shown in this heatmap.

**AI Summary**

Provider: ,
model:

Chat with Seqera AI

Export...

Copy prompt


Summarize plot

Sorted by sample

Clustered

Created with MultiQC

---

#### Software Versions

Software Versions lists versions of software tools extracted from file contents.

**AI Summary**

Provider: ,
model:

Chat with Seqera AI

 Copy table

| Software | Version |
| --- | --- |
| Cutadapt | `2.10` |
| FastQC | `0.11.9` |

**MultiQC v1.31** 
- Written by Phil Ewels,
available on GitHub.

This report uses Plotly,
jQuery,
jQuery UI,
Bootstrap and
FileSaver.js.

×

##### Plot Table Data

Select Column

Select Column

Please select two table columns.

Close

×

##### Regex Help

Toolbox search strings can behave as regular expressions (regexes). Click a button below to see an example of it in action. Try modifying them yourself in the text box.

`^` (start of string)
`$` (end of string)
`[]` (character choice)
`\d` (shorthand for `[0-9]`)
`\w` (shorthand for `[0-9a-zA-Z_]`)
`.` (any character)
`\.` (literal full stop)
`()` `|` (group / separator)
`*` (prev char 0 or more)
`+` (prev char 1 or more)
`?` (prev char 0 or 1)
`{}` (char num times)
`{,}` (count range)

```
samp_1
samp_1_edited
samp_2
samp_2_edited
samp_3
samp_3_edited
prepended_samp_1
tmp_samp_1_edited
tmpp_samp_1_edited
tmppp_samp_1_edited
#samp_1_edited.tmp
samp_11
samp_11111
```

See regex101.com for a more heavy duty testing suite.

Close
