## Supplementary Information (files S2-S7) for "A multi-omic, spatial, and whole-slide image dataset of lung neuroendocrine tumours from the lungNENomics cohort": S7.html

MultiQC Report


### Toggle navigation v1.31

Loading report..

- General Stats
- Space Ranger
  - Count - Summary stats
  - Count - Median genes
  - Count - Saturation plot
- Software Versions

Toolbox

##### MultiQC Toolbox

###### Apply Highlight Samples

+

Regex mode off
help
 Clear

###### Apply Rename Samples

+

Click here for bulk input.

Paste two columns of a tab-delimited table here (eg. from Excel).

First column should be the old name, second column the new name.

Add

Regex mode off
help
 Clear

###### Apply Show / Hide Samples

Hide matching samples

Show only matching samples

+

Regex mode off
help
 Clear

###### Explain with AI

Configure AI settings to get explanations of plots and data in this report.

AI Provider

Endpoint


Use the OpenAI API-style requests with a custom endpoint.

Model

API Key


Keys entered here will be stored in your browser's local storage.
See the docs.

Additional Payload


Any additional options passed in API request payload. Enter as a JSON object.

Context Window


The maximum number of tokens that can be processed in a single request


---

Anonymize samples off

###### Export Plots

- Images
- Data

px

px

Aspect ratio

PNG
SVG

Plot scaling

X

Download the raw data used to create the plots in this report below:

Format:

Tab-separated
Comma-separated
JSON

Note that additional data was saved in `multiqc_data` when this report was generated.

---

###### Choose Plots

 All
 None


---


   Download Plot Images

If you use plots from MultiQC in a publication or presentation, please cite:

> **MultiQC: Summarize analysis results for multiple tools and samples in a single report**  
> *Philip Ewels, Måns Magnusson, Sverker Lundin and Max Käller*  
> Bioinformatics (2016)  
> doi: 10.1093/bioinformatics/btw354  
> PMID: 27312411

Settings are automatically saved. You can also save named configurations below.

###### Save Settings

You can save the toolbox settings for this report to the browser or as a file.

 Save to Browser

 Save to File


---

###### Load Settings

Choose a saved report profile from the browser or load from a file:

[ select from browser ]

Load

 Delete

 Set default

 Clear default

Load from File

###### Tool Citations

Please remember to cite the tools that you use in your analysis.

To help with this, you can download publication details of the tools mentioned in this report:

List of DOIs

BibTeX file

###### About MultiQC

This report was generated using MultiQC, version 1.31

You can see a YouTube video describing how to use MultiQC reports here:
https://youtu.be/qPbIlO\_KWN0

For more information about MultiQC, including other videos and
extensive documentation, please visit http://multiqc.info

You can report bugs, suggest improvements and find the source code for MultiQC on GitHub:
https://github.com/MultiQC/MultiQC

MultiQC is published in Bioinformatics:

> **MultiQC: Summarize analysis results for multiple tools and samples in a single report**  
> *Philip Ewels, Måns Magnusson, Sverker Lundin and Max Käller*  
> Bioinformatics (2016)  
> doi: 10.1093/bioinformatics/btw354  
> PMID: 27312411

---

MultiQC is developed by:

A modular tool to aggregate results from bioinformatics analyses across many samples into a single report.

###### JavaScript Disabled

MultiQC reports use JavaScript for plots and toolbox functions. It looks like
you have JavaScript disabled in your web browser. Please note that many of the report
functions will not work as intended.

Loading report..

Report
generated on 2025-12-08, 15:33 CET
based on data in:
`/data/lungNENomics/files/Internal_Data_2023/Visium-already-on-tapes`

Summarize report

Copy report prompt

---

×
don't show again

**Welcome!** Not sure where to start?  
Watch a tutorial video
  *(6:06)*

**Report AI Summary**

More details...

Provider: ,
model: 

Chat with Seqera AI

#### General Statistics

**AI Summary**

Provider: ,
model:

Chat with Seqera AI

Table
 Export...

Copy prompt


Summarize plot

Created with MultiQC

Copy table

 Configure columns

 Sort by highlight

 Scatter plot

 Violin plot
Export as CSV...
Showing ^4^/_4_ rows and ^5^/_5_ columns.

Copy Prompt


Summarize table

| Sample Name | Reads | Spots Under Tissue | Reads In Spots | Avg Reads/Spot | Valid BC |
| --- | --- | --- | --- | --- | --- |
| LNEN071-IARC-A | 321.1M | 4009 | 89.5% | 80085 | 98.3% |
| LNEN084-IARC-B | 236.9M | 3542 | 86.1% | 66897 | 98.1% |
| LNEN107-IARC-C | 316.7M | 3582 | 92.4% | 88407 | 98.2% |
| LNEN206-IARC-D | 178.5M | 3062 | 80.4% | 58307 | 97.9% |

×

###### General Statistics: Columns

Uncheck the tick box to hide columns. Click and drag the handle on the left to change order. Table ID: `general_stats_table_table`

Show All
Show None

| Sort | Visible | Group | Column | Description | ID | Scale |
| --- | --- | --- | --- | --- | --- | --- |
| \|\| |  | Space Ranger: Space Ranger Count | Reads | Number of reads | `space_ranger_space_ranger_count-count_genstats_reads` | read\_count |
| \|\| |  | Space Ranger: Space Ranger Count | Spots Under Tissue | Number of Spots Under Tissue | `space_ranger_space_ranger_count-Count_spots_under_tissue` |  |
| \|\| |  | Space Ranger: Space Ranger Count | Reads In Spots | Fraction Reads in Spots Under Tissue | `space_ranger_space_ranger_count-Count_reads_in_spots` |  |
| \|\| |  | Space Ranger: Space Ranger Count | Avg Reads/Spot | Mean Reads per Spot | `space_ranger_space_ranger_count-Count_avg_reads_spot` |  |
| \|\| |  | Space Ranger: Space Ranger Count | Valid BC | Valid Barcodes | `space_ranger_space_ranger_count-Count_valid_bc` |  |

Close

#### Space Ranger

*spaceranger:*
 `1.3.0`

Tool to analyze 10x Genomics spatial transcriptomics data.*URL: https://support.10xgenomics.com/spatial-gene-expression/software/pipelines/latest/what-is-space-ranger*

##### Count - Summary stats

Summary QC metrics from Space Ranger count

**AI Summary**

Provider: ,
model:

Chat with Seqera AI

Table
 Export...

Copy prompt


Summarize plot

Created with MultiQC

Copy table

 Configure columns

 Sort by highlight

 Scatter plot

 Violin plot
Export as CSV...
Showing ^4^/_4_ rows and ^7^/_14_ columns.

Copy Prompt


Summarize table

| Sample Name | Reads | Spots Under Tissue | Reads In Spots | Avg Reads/Spot | Median UMI/Spot | Median Genes/Spot | Genes Detected | Valid BC | Valid UMI | Saturation | Q30 BC | Q30 UMI | Reads Mapped | Confident Reads |
| --- | --- | --- | --- | --- | --- | --- | --- | --- | --- | --- | --- | --- | --- | --- |
| LNEN071-IARC-A | 321.1M | 4009 | 89.5% | 80085 | 21769.0 | 6834 | 17887 | 98.3% | 100.0% | 63.5% | 96.1% | 96.1% | 98.5% | 93.1% |
| LNEN084-IARC-B | 236.9M | 3542 | 86.1% | 66897 | 14197.0 | 5318 | 17835 | 98.1% | 100.0% | 69.5% | 96.3% | 96.3% | 97.7% | 96.4% |
| LNEN107-IARC-C | 316.7M | 3582 | 92.4% | 88407 | 13096.0 | 5078 | 17830 | 98.2% | 100.0% | 82.2% | 96.0% | 96.1% | 97.5% | 95.7% |
| LNEN206-IARC-D | 178.5M | 3062 | 80.4% | 58307 | 3834.0 | 1876 | 17782 | 97.9% | 100.0% | 89.3% | 96.1% | 96.2% | 97.0% | 93.3% |

×

###### Space Ranger: Count: Summary stats: Columns

Uncheck the tick box to hide columns. Click and drag the handle on the left to change order. Table ID: `spaceranger-count-stats_table`

Show All
Show None

| Sort | Visible | Group | Column | Description | ID | Scale |
| --- | --- | --- | --- | --- | --- | --- |
| \|\| |  |  | Reads | Number of reads | `space_ranger_count-count_data_reads` | read\_count |
| \|\| |  | Space Ranger Count | Spots Under Tissue | Number of Spots Under Tissue | `space_ranger_count-Count_spots_under_tissue` |  |
| \|\| |  | Space Ranger Count | Reads In Spots | Fraction Reads in Spots Under Tissue | `space_ranger_count-Count_reads_in_spots` |  |
| \|\| |  | Space Ranger Count | Avg Reads/Spot | Mean Reads per Spot | `space_ranger_count-Count_avg_reads_spot` |  |
| \|\| |  | Space Ranger Count | Median UMI/Spot | Median UMI Counts per Spot | `space_ranger_count-Count_median_umi_spot` |  |
| \|\| |  | Space Ranger Count | Median Genes/Spot | Median Genes per Spot | `space_ranger_count-Count_median_genes_spot` |  |
| \|\| |  | Space Ranger Count | Genes Detected | Genes Detected | `space_ranger_count-Count_genes_detected` |  |
| \|\| |  | Space Ranger Count | Valid BC | Valid Barcodes | `space_ranger_count-Count_valid_bc` |  |
| \|\| |  | Space Ranger Count | Valid UMI | Valid UMIs | `space_ranger_count-Count_valid_umi` |  |
| \|\| |  | Space Ranger Count | Saturation | Sequencing Saturation | `space_ranger_count-Count_saturation` |  |
| \|\| |  | Space Ranger Count | Q30 BC | Q30 Bases in Barcode | `space_ranger_count-Count_Q30_bc` |  |
| \|\| |  | Space Ranger Count | Q30 UMI | Q30 Bases in UMI | `space_ranger_count-Count_Q30_UMI` |  |
| \|\| |  | Space Ranger Count | Reads Mapped | Reads Mapped to Probe Set | `space_ranger_count-Count_reads_mapped` |  |
| \|\| |  | Space Ranger Count | Confident Reads | Reads Mapped Confidently to Probe Set | `space_ranger_count-Count_confident_reads` |  |

Close

---

##### Count - Median genes Help

Median gene counts per spot

This plot shows the Median Genes per Spot as a function of downsampled sequencing depth in mean reads per spot, up to the observed sequencing depth. The slope of the curve near the endpoint can be interpreted as an upper bound to the benefit to be gained from increasing the sequencing depth beyond this point.

**AI Summary**

Provider: ,
model:

Chat with Seqera AI

Export...

Copy prompt


Summarize plot

Created with MultiQC

---

##### Count - Saturation plot Help

Sequencing saturation

This plot shows the Sequencing Saturation metric as a function of downsampled sequencing depth (measured in mean reads per spot), up to the observed sequencing depth. Sequencing Saturation is a measure of the observed library complexity, and approaches 1.0 (100%) when all converted probe ligation products have been sequenced. The slope of the curve near the endpoint can be interpreted as an upper bound to the benefit to be gained from increasing the sequencing depth beyond this point. The dotted line is drawn at a value reasonably approximating the saturation point.

**AI Summary**

Provider: ,
model:

Chat with Seqera AI

Export...

Copy prompt


Summarize plot

Created with MultiQC

---

#### Software Versions

Software Versions lists versions of software tools extracted from file contents.

**AI Summary**

Provider: ,
model:

Chat with Seqera AI

 Copy table

| Group | Software | Version |
| --- | --- | --- |
| Space Ranger | spaceranger | `1.3.0` |

**MultiQC v1.31** 
- Written by Phil Ewels,
available on GitHub.

This report uses Plotly,
jQuery,
jQuery UI,
Bootstrap and
FileSaver.js.

×

##### Plot Table Data

Select Column

Select Column

Please select two table columns.

Close

×

##### Regex Help

Toolbox search strings can behave as regular expressions (regexes). Click a button below to see an example of it in action. Try modifying them yourself in the text box.

`^` (start of string)
`$` (end of string)
`[]` (character choice)
`\d` (shorthand for `[0-9]`)
`\w` (shorthand for `[0-9a-zA-Z_]`)
`.` (any character)
`\.` (literal full stop)
`()` `|` (group / separator)
`*` (prev char 0 or more)
`+` (prev char 1 or more)
`?` (prev char 0 or 1)
`{}` (char num times)
`{,}` (count range)

```
samp_1
samp_1_edited
samp_2
samp_2_edited
samp_3
samp_3_edited
prepended_samp_1
tmp_samp_1_edited
tmpp_samp_1_edited
tmppp_samp_1_edited
#samp_1_edited.tmp
samp_11
samp_11111
```

See regex101.com for a more heavy duty testing suite.

Close
