## Supplementary figure for "A multi-omic, spatial, and whole-slide image dataset of lung neuroendocrine tumours from the lungNENomics cohort"

Supplementary Figures

CONTENTS

Supplementary Figure S1

Supplementary Figure S2

Supplementary Figure S3

Supplementary Figure S4

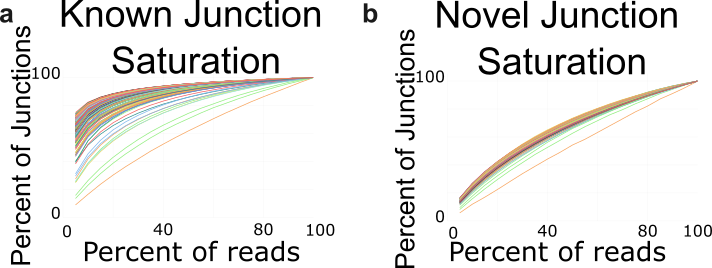
**Supplementary Figure S1**. **Junction Saturation in RNA-seq** (a) Number of known junctions identified by software STAR in a RNA-seq subsample as a function of the percentage of reads in the subsample. (b) Number of novel junctions identified by STAR in a RNA-seq subsample as a function of the percentage of reads in the subsample.

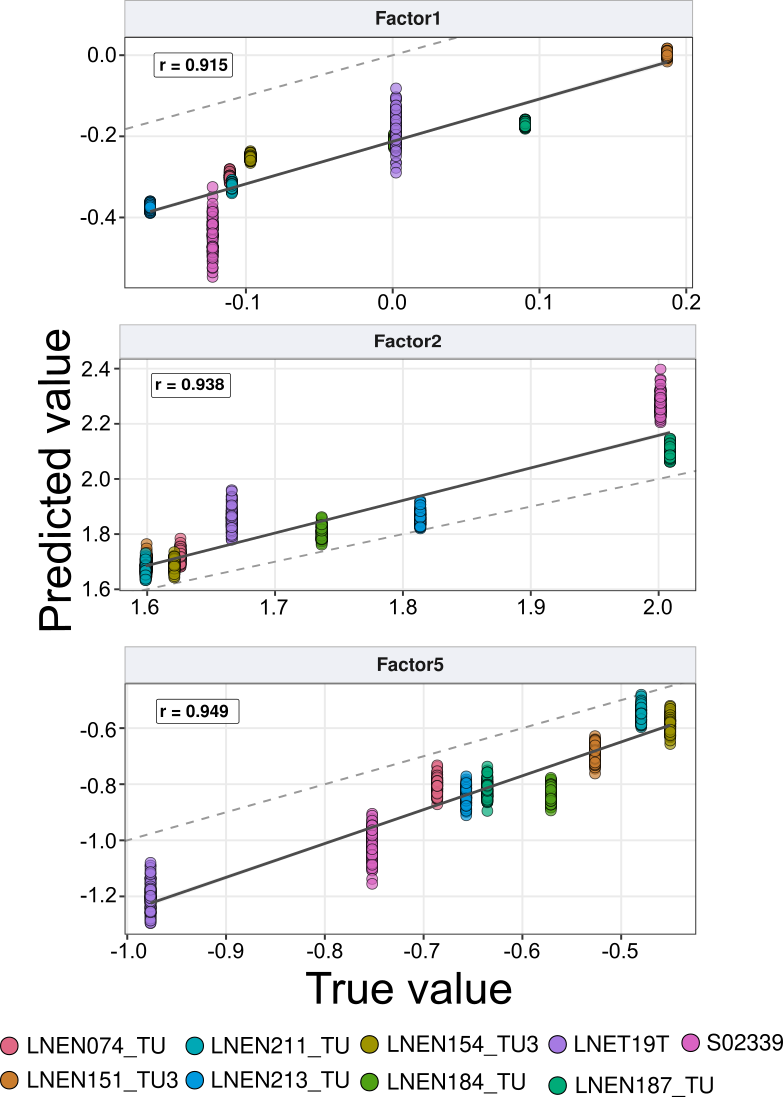

**Supplementary Figure S2**. **Concordance between original and downsampling projected MOFA2 latent factor values.** Scatter plot shows the true (original full depth) versus predicted (downsampled) MOFA2 latent factor values for Factor1, Factor2, and Factor5 across 9 validation samples, each simulated across 100 downsampling iterations mimicking LNEN079_TU library size. Each dot represents one iteration per sample, coloured by sample. The solid line indicates linear regression fit; dashed line indicates perfect concordance (r=1). Pearson correlation coefficients (r) are shown per factor at the top left.

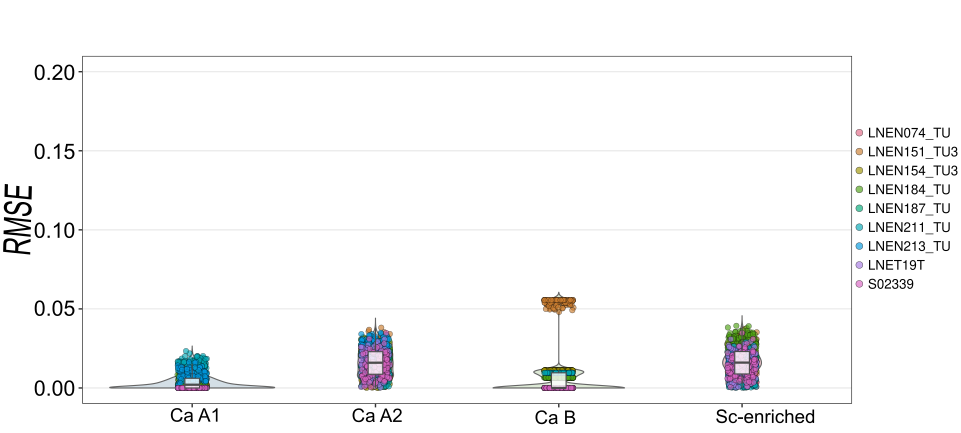

**Supplementary Figure S3**. **Robustness of archetype proportions under simulated low sequencing depth.** Root Mean Square error (RMSE) between original (full sequencing depth) and downsampling-projected archetype proportions across 100 simulated iterations per sample, shown for each of the four archetypes (CA A1, Ca A2, Ca B, Sc-enriched). Each dot represents one iteration from one of the 9 validation samples, coloured by sample.

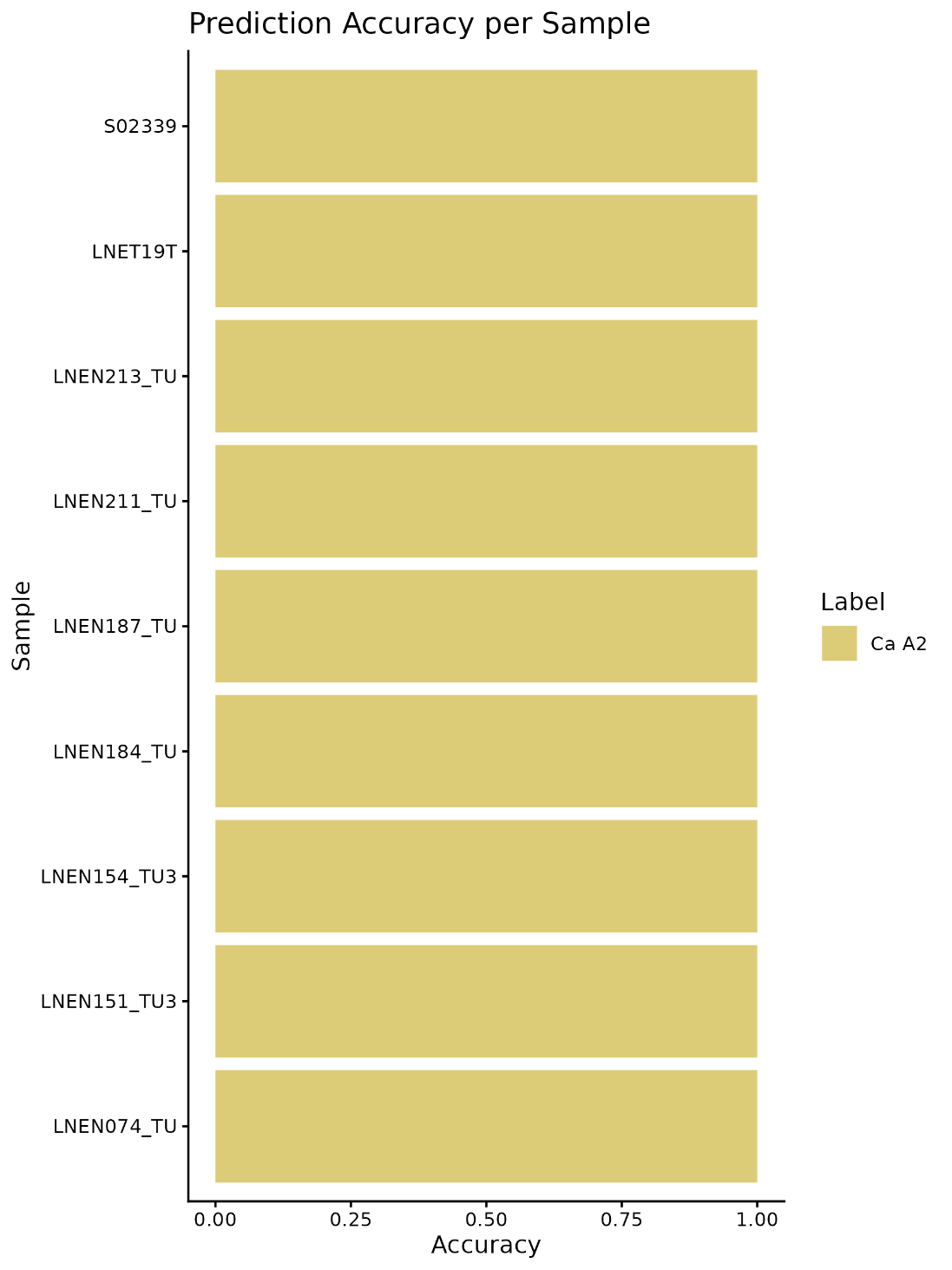

**Supplementary Figure S4**. **Prediction Accuracy of downsampled samples.** The proportion of predicted archetype labels across repeated iterations.
