## Supplementary information File S1 for "A multi-omic, spatial, and whole-slide image dataset of lung neuroendocrine tumours from the lungNENomics cohort"

**DATA ACCESS AGREEMENT**

**Reference Number: DTA-2026-XXX**

**THIS Data Access Agreement (“Agreement”)** enters into full force and effect as of the date of its full signature **(“Effective Date”)**

**BETWEEN:**

**The International Agency for Research on Cancer, World Health Organization** of 25 avenue Tony Garnier, CS 90627, 69366 Lyon Cedex 07, France **(“IARC/WHO”),** with Dr Matthieu Foll as IARC/WHO’s Principal Investigator hereunder; and

**[NAME AND ADRESS OF RECEIVING INSTITUTE]** (the “**Receiving Institute**”), with [TITLE AND NAME OF RECEIVING PI] as Receiving Institute’s Principal Investigator hereunder;

(each a “**Party**” and collectively the “**Parties**”).

**The Parties AGREE** **AS FOLLOWS**:

1. This Agreement will start on the Effective Date stated above and, unless earlier terminated, will end on XXX (“**End Date**”).
2. The Receiving Institute has requested access to the following data (hereinafter “Data”):

The data hosted on the European Genome-phenome Archive (EGA) under the study accession number EGAS00001005979. This study contains raw multi-omics data generated as part of the lungNENomics project, a multi-omics study aimed at deep molecular profiling of lung neuroendocrine tumours, including typical carcinoids (grade 1) and atypical carcinoids (grade 2). The study includes whole-genome sequencing (WGS), RNA sequencing (RNA-seq), and Infinium MethylationEPIC array data from samples in the lungNENomics cohort.

This study includes four datasets accession numbers:

EGAD00001015672: Annotated germline variant calls derived from WGS data for 72 individuals.

EGAD00001015524: Raw RNA-seq data for 179 individuals.

EGAD00001015520: Raw WGS data for 72 individuals.

EGAD00010002480: Raw DNA methylation array data for 181 individuals.

for the purpose of the project entitled:

[PROJECT TITLE] (the “**Research Project**”);

The Research Project is further described in **Appendix 1**.

3. IARC/WHO will authorize access to the Data via the EGA portal to the Receiving Institute upon receipt of the fully-executed copy of this Agreement acknowledging and agreeing to its terms.
4. **Annex 1** of this Agreement describes the obligations of the Receiving Institute regarding the security it will apply to the Data to which it is granted access under this Agreement. Annex 1 (and any other annexes or appendices) forms an integral part of this Agreement and is legally binding between the Parties.

**Authorized use of the Data:**

5. The Data are made available to the Receiving Institute under this Agreement solely for non-profit research, and solely in connection with and for the purpose of the Research Project.
6. Other than for and within the purpose of the Research Project, as described in Appendix 1, the Data will not be further transferred, distributed to third parties or otherwise used without IARC/WHO’s prior written consent.
7. The Data will be used only and solely by the Receiving Institute and Receiving Institute’s authorized personnel, under the responsibility and supervision of the Receiving Institute’s Principal Investigator, for the purposes hereof exclusively and under no less stringent obligations than as provided for in this Agreement.
8. The Receiving Institute will not seek to reverse engineer or de-anonymize the Data in any way whatsoever and will comply with any applicable ethical requirements. The Receiving Institute further represents and warrants that the use of the Data will not violate any acts, laws, by-laws, rules and regulations applicable to the Data.

**Confidentiality:**

9. The Receiving Institute agrees to keep the Data in confidence, except for Data that: (a) are publicly known, or available from other sources which are not under a confidentiality obligation to the source; (b) have been made available by its owners without a confidentiality obligation; (c) are otherwise already known by or available to the Receiving Institute without a confidentiality obligation; or (d) are required to be disclosed by operation of law, provided that the Receiving Institute immediately so notifies IARC/WHO in writing and provides adequate opportunity for IARC/WHO to object to, or restrict, such disclosure or request confidential treatment thereof.

**Intellectual property rights and ownership:**

10. Except for the rights explicitly granted hereunder, nothing contained in this Agreement is construed as conveying any rights under any patents or other intellectual property which either Party may have as of the date of this Agreement or may hereafter obtain, independently of and without reference to the Research Project or the Data.
11. IARC/WHO retains ownership and/or custody of the Data as applicable and has the unrestricted right to use, disclose or transfer the Data to any third parties for any other purposes. The Receiving Institute acknowledges and agrees that nothing contained in this Agreement is deemed to grant to the Receiving Institute any intellectual property rights in any of the Data to which it is granted access hereunder.

**Warranties and liability:**

12. IARC/WHO makes no warranty of the fitness of the Data for any particular purpose or any other warranty, either express or implied.
13. The Receiving Institute agrees that, except as may explicitly be provided in this Agreement, IARC/WHO has no control over the Receiving Institute’s use of the Data hereunder. Consequently, the Receiving Institute agrees that IARC/WHO shall not be liable for such use, or any loss, claim or damages which may arise from or in connection with such use.

**Miscellaneous:**

14. Nothing contained in this Agreement will be construed as a waiver of any of the privileges and immunities enjoyed by IARC/WHO, as part of the World Health Organization (WHO) and the United Nations system, under national or international law, and/or as submitting IARC/WHO to any national law or court jurisdiction.
15. Any dispute relating to the interpretation or application of this Agreement will, unless amicably settled, be subject to conciliation. In the event of failure of the latter, the dispute will be settled by arbitration. The arbitration will be conducted in accordance with the modalities to be agreed upon by the Parties or, in the absence of agreement, with the rules of arbitration of the International Chamber of Commerce. The Parties will accept the arbitral award as final.
16. This Agreement may be amended only by written agreement duly signed by the authorized representatives of the Parties. Neither Party will assign, transfer, or deal in any other manner with its rights and obligations under this Agreement without the express prior written consent of the other Party.
17. IARC/WHO reserves the right to terminate this Agreement upon 30 days’ written notice to the Receiving Institute.
18. Upon expiry or earlier termination of this Agreement, the Receiving Institute will immediately cease any use, and securely dispose of the Data, delete the Data or otherwise in accordance with IARC/WHO’s instructions.
19. This Agreement will in no way be construed as creating the relationship of principal and agent, of partnership in law or of joint venture as between the Parties or any other person involved in the Research Project.
20. The electronic signature via DocuSign will have the same validity, legality and enforceability as an original signature. Each of the Parties receives a fully signed copy of this Agreement. The delivery of such fully signed copy via e-mail or DocuSign will have the same validity as the delivery of an original.

[*Signatures and acknowledgements follow on the next page*]

This Agreement is duly signed on behalf of the Parties as follows:

| Signed for and on behalf of the Receiving Institute: | Signed for and on behalf of IARC/WHO: |
| --- | --- |
| ________________________________ | ________________________________ |
| The Receiving Institute’s Authorized Official | IARC/WHO’s Authorized Official |
| Name: | Name: |
| Title: | Title: Director of Administration & Finance |
| Date: Read and acknowledged by the Principal Investigator of the Receiving Institute: ________________________________; Name:; Title:; Email:; Date: | Date: Read and acknowledged by the Principal Investigator of IARC/WHO: ________________________________; Name:; Title: Principal Investigator; Email:; Date: |

**ANNEX 1**

**SECURITY TO BE APPLIED BY the Receiving Institute TO THE DATA**

The following security measures and controls must be respected at all times:

• Access Control: Physical and logical access controls must be established in order to protect the Data at all times. The Data should be stored in a secure location requiring either badge or key access to its physical location. Logical access will be controlled with Access Controls Lists (ACL), username and strong password combinations and file and share level permissions.

• User Access: Access to the Data will only be granted by the Principal Investigator (PI). A central log of users having access to the Data should be maintained.

• Passwords: All passwords should be strong in nature. Passwords must never be written down nor shared. Usernames and passwords should be, if possible centrally managed.

• Cryptography: When the data is of human origin it should be encrypted when stored on laptops or removal storage devices such as external disks.

• Virus Protection: All IT equipment storing or accessing the Data should have up to date antivirus protection.

• Operating System Management: All IT equipment storing or accessing the data must be updated automatically on a regular basis with operating system and security patches in order to avoid potential security breaches. Ideally, this should be managed centrally.

• Backup: The Data should be backed up on a regular basis and stored in a separate location to the original data in order to allow the recovery of the data after a major incident.

• Logging: Logging of access to the Data should be put in place in order to allow a clear audit trail to be maintained of access and modifications made by each authorized User.

• Network Security: The network where the data is stored should be secured to avoid unauthorized access to the information. Network segregation and firewalls should implemented to increase the safety of the data.

**APPENDIX 1**

**DESCRIPTION OF THE RESEARCH PROJECT**
